# Modeling the journey as well as the destination: a control theory account of rotational navigation

**DOI:** 10.64898/2026.06.02.729319

**Authors:** Yinqi Huang, Abhilasha Vishwanath, Yu Karen Du, Matthew F Watson, Osama Asiri, Karlee Dakin, Donette Markham, Arne D Ekstrom, Robert C Wilson

## Abstract

Navigation requires estimating heading and transforming these estimates into actions. Prior models explain how self-motion and landmark cues are combined into heading estimates, but less is known about how these estimates are iteratively transformed into motor commands to reach a goal. Here, we hypothesized that navigation operates as a closed-loop process in which ongoing movement is updated by sensory prediction errors. To test this hypothesis, participants performed a goal-directed rotation task in virtual reality. On select trials, visual landmarks were shifted during movement, inducing a prediction error between the heading expected from self-motion estimates and the heading observed from the shifted landmarks. In parallel, we developed a closed-loop model of turning behavior that represents heading and angular velocity as jointly estimated states over time. This model accounts not only for final position—the destination—but also for the movement dynamics that produce it—the journey. The model predicts that landmark-induced visual prediction errors should produce rapid corrective changes in movement. Participant turning behavior qualitatively paralleled these model dynamics: acceleration changed after visual feedback, with larger landmark mismatches producing larger corrective responses. Together, these findings suggest that naturalistic movement depends on continuously transforming heading estimates into motor command through closed-loop control.

## 1 Introduction

Goal-directed behavior requires more than knowing where the body is relative to a target. When reaching for a coffee mug, for example, visual and proprioceptive information are used not only to estimate the hand’s current position, but also to update motor commands as the hand moves toward the mug. Real-world navigation poses an analogous problem: reaching a remembered destination requires iteratively updating both the estimated heading and the motor commands used to guide movement. For example, when walking toward a supermarket, a mismatch between the expected and observed position of the store may indicate that one has walked too far west, requiring an online correction of both the internal heading estimate and the trajectory needed to reach the goal. This behavior depends in part on path integration: the process of maintaining an internal estimate of heading by integrating idiothetic selfmotion cues over time [1–3]. These cues include optic flow, vestibular, proprioceptive, and motor-related signals, such as efference copies of motor commands [4–9]. Stable landmarks, such as stores, can also recalibrate internal estimates of heading, a process often referred to as landmark navigation [10, 11].

In everyday navigation, humans and other animals navigate using a combination of path integration and external landmarks. A large body of work has investigated how these cues are combined to form internal estimates of heading and position. This cue-combination process is often formalized as an observer model that updates its estimate by weighting incoming sensory information according to its reliability, yielding a possibly optimal estimate of the current state [12–18]. Despite differences in implementation, such observer models share a common limitation: they primarily explain endpoint estimates. As a result, they do not specify how an internal estimate of heading is iteratively transformed into motor commands needed to reach a navigational goal. Experimentally, this limitation manifests in a focus on the final response — the destination — while ignoring the control dynamics to get there — the journey. Recent work has begun to address this gap by framing navigation as control under spatial uncertainty, accounting for both trajectories and endpoint variability using a probabilistic model of an agent’s belief about its location and orientation [19]. In that framework, control is defined at the level of an optimal policy that maps belief states to actions. It remains unclear, however, how sensory prediction errors are translated into moment-by-moment changes in motor output, such as acceleration. This omission matters because movement trajectories can reveal control dynamics that is not available from endpoint measures alone.

We address this gap by testing whether angular path integration can be modeled as a problem of state estimation and feedback control. The approach is motivated by control-theoretic models of motor behavior, in which actions such as reaching or smooth pursuit are continuously updated using sensory feedback, with motor corrections applied to reduce deviations from the goal [20–22]. Optimal feedback control theories further propose that movements are produced by control policies that minimize task-relevant costs while allowing variability in task-irrelevant dimensions [23–26]. These models provide a mechanistic model to represent feedback control, but they often assume that the goal state is already known. Navigation, in contrast, often re-estimates the states by integrating self-motion and external sensory cues throughout movement.

Despite their complementary insights, these two lines of research have largely developed in isolation and therefore provide incomplete accounts of goal-directed navigation. Navigation research emphasizes how sensory cues are combined to estimate heading, whereas motor-control research emphasizes how control policies generate actions given state estimates and a goal. Recent work on human walking has shown that movement dynamics contain information beyond endpoint accuracy [27]. A useful model of rotational navigation should therefore explain not only where people stop, but also how heading, velocity, and acceleration evolve as they move toward a goal.

In this paper, we test whether a simple control-theoretic model is sufficient to generate the qualitative patterns observed in human rotational data. In this closed-loop model, internal estimates of heading and angular velocity are updated by prediction errors and then used to generate motor commands through a control law. Here, we do not treat the model as a quantitative fit to human trajectories. Instead, we use it as an existence proof of architectural sufficiency: a system combining state estimation, prediction error from observation, and closed-loop feedback control can generate velocity and trajectory profiles that account for endpoint errors and feedback-induced acceleration corrections observed in human turning behavior. This model provides a testable baseline for future quantitative model fitting.

## 2 Results

105 participants performed an angular path integration task (the Rotation Task [15]) in immersive virtual reality. 34 participants performed one session of the task, 71 participants completed two sessions separated by 3–14 days for a total of 176 behavioral sessions. In each trial, participants first encoded an angle by rotating to a specified heading and then attempted to reproduce that angle from memory by returning to the original heading. The key experimental manipulation was whether visual landmark feedback was provided during reproduction and, when present, the extent to which the landmarks were offset relative to their correct location. In the No Feedback condition, participants viewed a blank screen during reproduction and relied solely on self-motion cues (Figure 1b). In the Feedback condition, the virtual room was briefly visible for 300 ms at time *t*_*fb*_, at jitter angle *fb*, with the room rotated by a jitter angle *j* relative to the participant’s true heading (Figure 1c). In the majority of the Feedback trials (probability *ρ* = 0.7), the jitter angle was drawn from a Gaussian centered on the true heading (*σ* = 30°), providing approximately consistent visual information. In the remaining trials (probability 1 − *ρ* = 0.3), the jitter angle was drawn from a uniform distribution, which introduces large and unpredictable mismatches between the visual scene and the participant’s internal heading estimate. For the 34 participants in the one-session dataset, jitter angles were sampled uniformly between −100° and +100° (−5*π/*9 to + 5*π/*9 rad), whereas for the 71 participants in the two-session dataset, jitter angles were sampled uniformly between −180° to +180° (−*π* to + *π* rad).

**Fig. 1.**
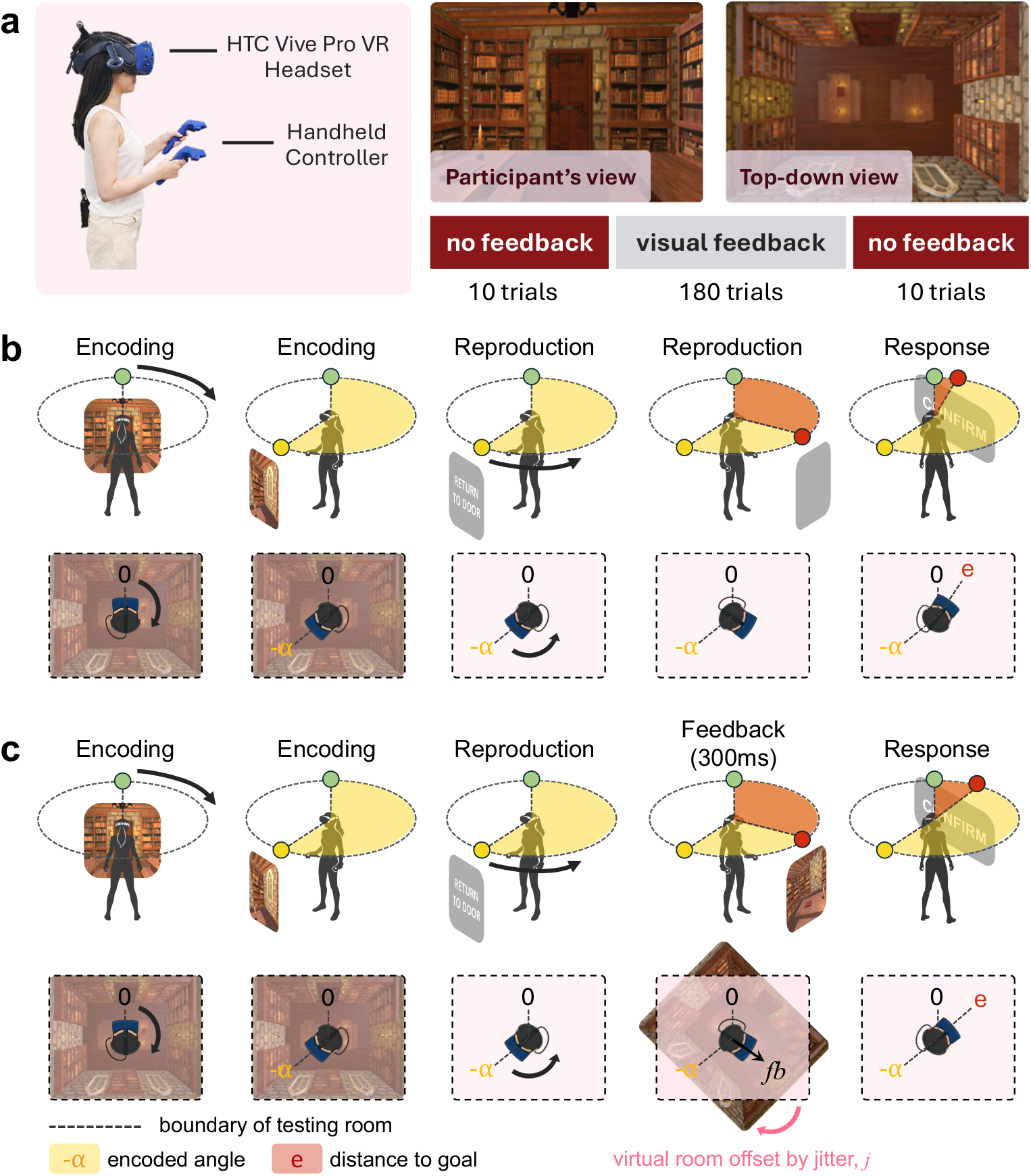
Experimental design of the Rotation Task. **(a) Experimental setup**. An author of this paper wore an HTC Vive Pro head-mounted display and held two handheld controllers while turning in a square virtual room. The individual shown in the photograph is an author of this paper. The experiment consisted of three blocks: 10 no-feedback trials, 180 feedback trials, and 10 no-feedback trials. First-person and top-down views of the virtual environment are shown. **(b) No Feedback condition**. For illustration, the figure depicts a clockwise turn (i.e., −*α*). Participants began at the door location (heading 0), were guided to rotate to an encoded angle −*α*, and then turned back by their estimated traveled angle 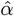 toward the remembered door location without visual feedback. The goal was to rotate in the opposite direction and return to the original heading (0). **(c) Feedback condition**. The procedure is identical except that, at feedback onset, the virtual room was offset by a jitter angle *j*. The visual scene was rotated by *j* relative to its true orientation, generating a visual feedback signal *fb*. Participants updated their internal estimate 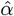 and adjusted their turning behavior to compensate for this prediction error. The distance to goal *e* is defined as the difference between the reproduced angle and the encoded angle.

### 2.1 Reproduced angle reveals cue combination and competition during rotation

We first asked whether the current study reproduced endpoint behavior previously reported in [15]. In the No Feedback condition, Figure 2a–c shows that distance to goal varies as a function of encoded angle, with the slope of this relationship varying across individuals: some participants systematically underestimate small goal angles (positive slope), others underestimate large goal angles (negative slope), and some show approximately zero slope. This pattern is consistent with previous results from the same task.

**Fig. 2.**
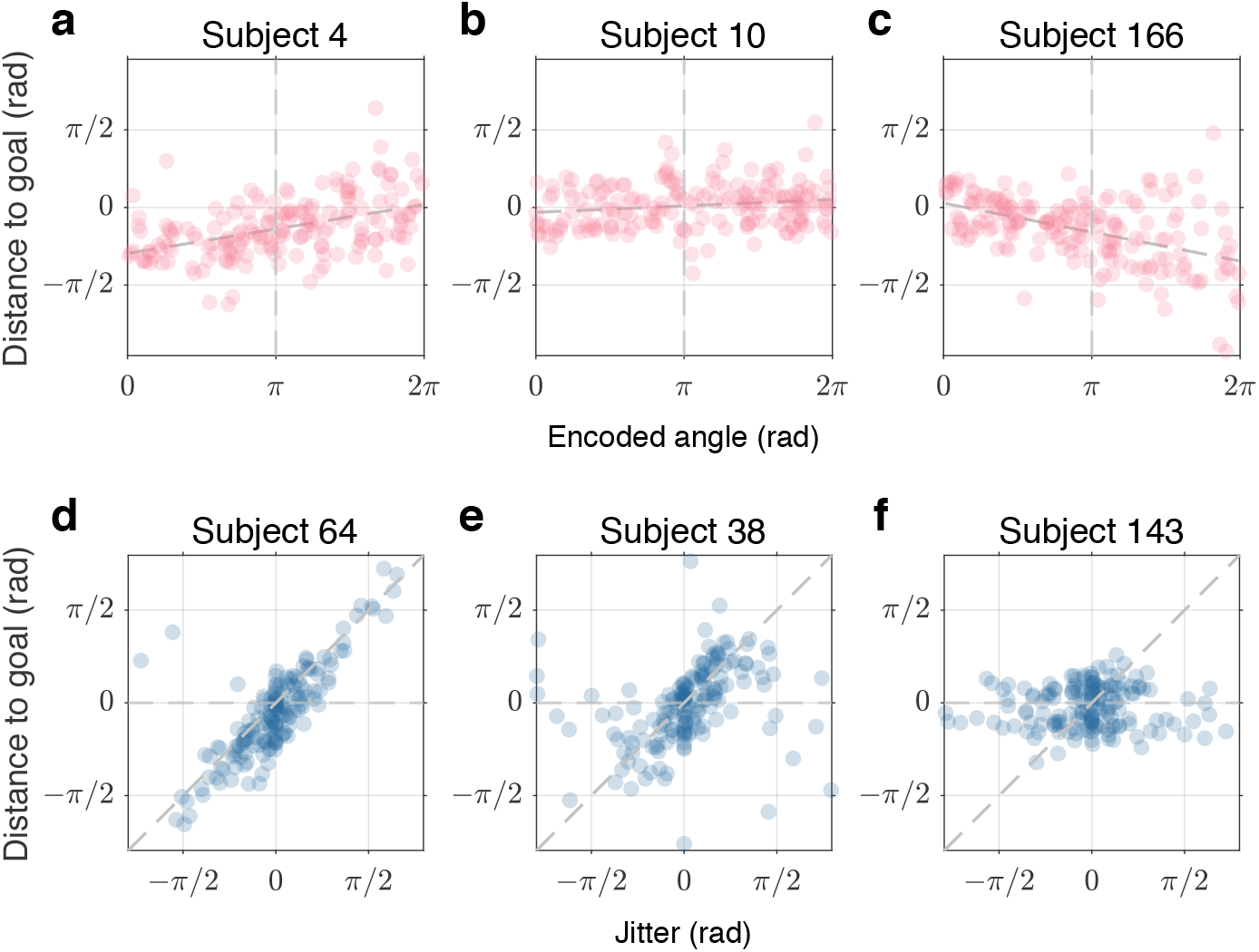
Individual differences in angular path integration and landmark navigation. Data from six example participants, with each point representing a single trial. **(a–c) Encoded angle versus distance to goal**. Distance to goal varies as a function of encoded angle, and the slope of this relationship differs across individuals. Some participants showed a positive slope, indicating systematic underestimation of small goal angles; others showed a negative slope, indicating underestimation of large goal angles; and some showed an approximately zero slope. The dashed line represents a linear fit (least-squares regression) to the data, summarizing the dependence of distance to goal on encoded angle for each participant. **(d–f) Visual jitter versus distance to goal**. Distance to goal varies as a function of induced visual jitter. The slope of this relationship differs across individuals, with some participants showing stronger dependence on visual landmarks than others. This variability is consistent with differences in the weighting of visual feedback during cue combination reported previously in [15].

Next, we examined the influence of visual feedback in the Feedback condition. Figure 2d–f shows that response error systematically varied with room jitter for many participants, indicating that visual landmarks influenced estimates of the original heading. At the same time, the strength of this relationship differed across individuals: some participants showed substantial changes in response error as jitter increased, whereas others were insensitive to visual feedback. These findings are consistent with previous work showing individual differences in the weighting of visual and self-motion cues during navigation [15]. Prior observer model explained such variances in terms of differences in how navigators weighted the reliability of visual feedback when estimating heading. However, these models focused primarily on endpoint estimates and did not address how sensory feedback is transformed into the moment-to-moment motor commands. This limitation motivates our next step: a control-theoretic account of the Rotation Task that links heading estimation to continuous motor control.

### 2.2 A control-theoretic account for both reproduced angle and turning dynamics

To ask whether closed-loop control could generate the observed movement dynamics in participants, we modeled angular path integration as a process combining state estimation and feedback control. In this model, the navigator maintains an internal estimate of heading and angular velocity, and continuously updates this estimate based on internal dynamics and sensory observations. Motor commands are generated to minimize the discrepancy between the estimated state and the goal. The model consists of three interacting components: the real-world state that the system is trying to control, an observer that estimates the current state, and a controller that generates control signals to drive the system toward the goal (Figure 3). The model contains several free parameters, so the present data do not uniquely identify a single parameterization. We therefore use the model for qualitative pattern comparison rather than as a quantitative fit to individual trajectories. However, the simulated trajectories, including velocity and acceleration profiles, remain within the empirical range observed in the human data, such that the simulation operates at a realistic rather than arbitrary or biologically implausible scale.

**Fig. 3.**
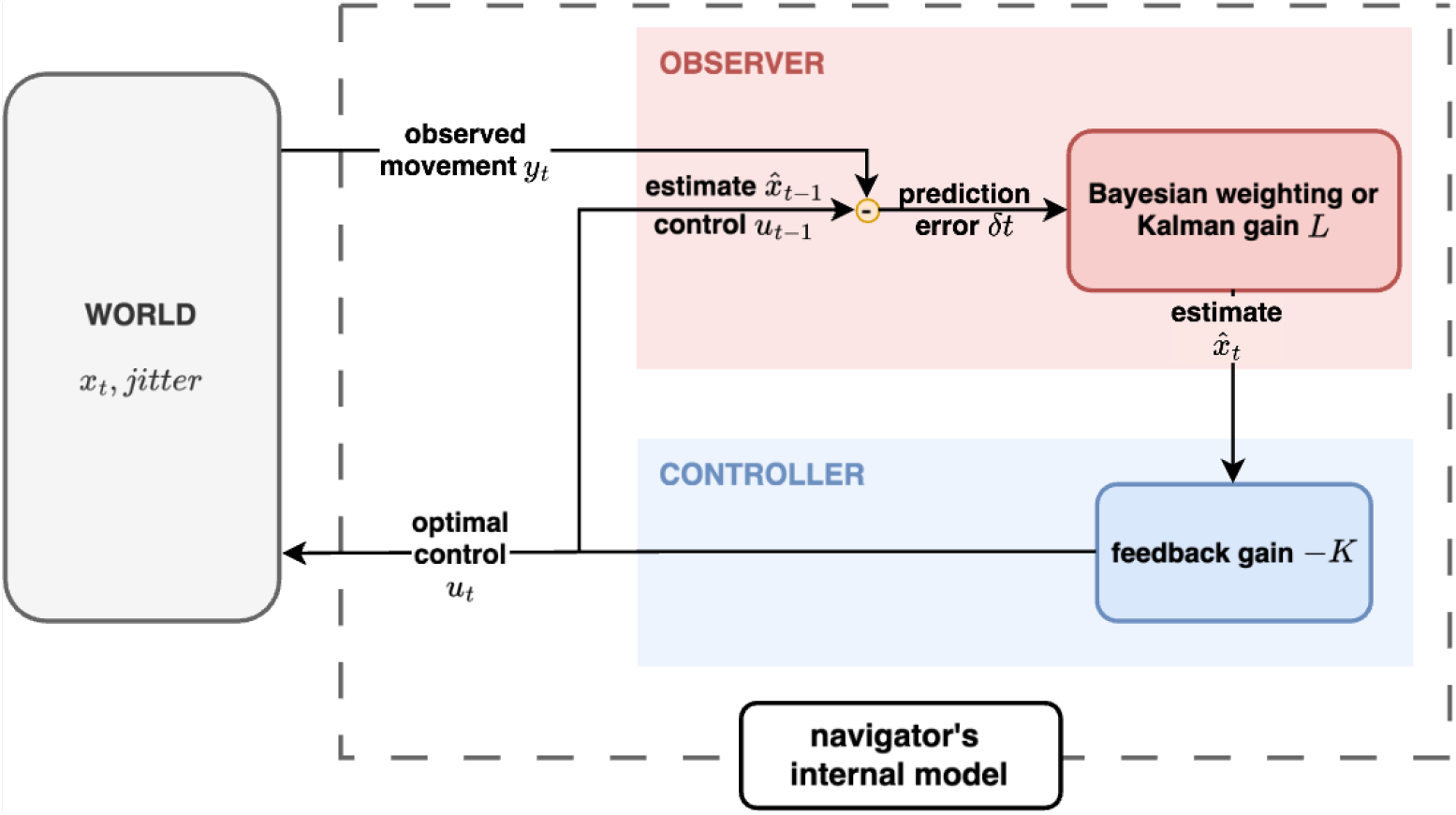
Control theory model of the Rotation Task. The navigator’s internal estimate of heading and angular velocity is modeled as part of a closed-loop system consisting of an observer that estimates the current state and a controller that generates motor commands to minimize deviation from the goal.

#### World: Real-world state to be controlled

The real-world state describes the physical dynamics of turning. Let *h*_*t*_ denote heading and *v*_*t*_ denote angular velocity. In the Rotation Task, participants generate a control input *u*_*t*_, which manifests as an angular acceleration. We model participant turning behavior as a simple dynamical system in which heading evolves as the time integral of angular velocity, while angular velocity is driven by acceleration (control input) and opposed by a velocity-proportional resistance term (capturing effects such as viscous damping and floor friction). For simplicity, we neglect the second-order acceleration contribution to the heading update and approximate the dynamics in discrete time using first-order Taylor expansion:

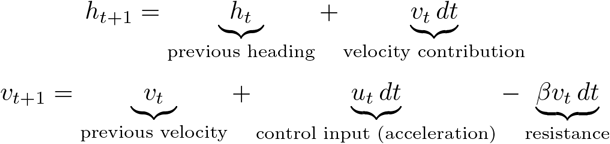

where *dt* is the time step and *β* ≥ 0 is a resistance coefficient.

We now express these dynamics in state-space form. Turning behavior is quantified by two variables: the heading angle *h*_*t*_ and its velocity *v*_*t*_, which correspond to the data measured in the experiment. Together, they form the state vector

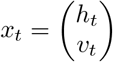

In the Rotation Task, the participant’s goal is to move the state from the encoded angle −*α* to the door at angle 0, with zero angular velocity at both the start and the end (Figure 1). Thus, the controller must move from the initial state

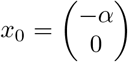

to the goal state

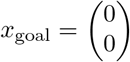

To reach the goal, participants generate a control signal *u*_*t*_, which updates the state according to

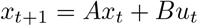

The matrices *A* and *B* follow directly from the discretized physical dynamics above. In particular, the state transition matrix *A* is written as

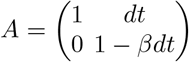

which reflects that heading integrates velocity, and velocity decays over time due to resistance. And the control input matrix *B* is

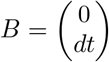

which updates angular velocity.

#### Observer: Estimated current state

The observer describes participant’s internal estimation of the real-world states. Participants do not observe the true state of the world *x*_*t*_ directly. Instead, they must infer their heading and angular velocity by integrating self-motion cues over time and combining these internal estimates with visual and other sensory cues when they are available. We write this estimated state as

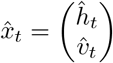

which evolves over time according to

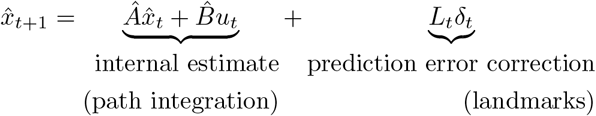

where *Â* is the participant’s internal estimate of the dynamics matrix, 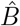 is the participant’s estimate of the effect of the control signal. *L*_*t*_ is the gain applied to sensory prediction error, and *δ*_*t*_ is the mismatch between predicted and observed sensory input. In general, *Â* ≠ *A* and 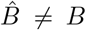, which captures the possibility that participants’ internal model of the dynamics differs from the real-world dynamics [28].

We define these matrices as

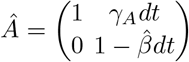

and

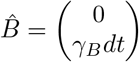

The gain parameters *γ*_*A*_ and *γ*_*B*_ capture systematic mismatches between the participant’s internal estimate and the world dynamics: *γ*_*A*_ scales the internal estimate of velocity, whereas *γ*_*B*_ scales the internal estimate of acceleration. This formulation is consistent with prior work showing that path integration errors accumulate with self-motion [8, 29]. For simplicity, we assume that the participant’s estimate of resistance matches the true resistance, that is, 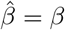. While this assumption could be relaxed, it simplifies the analysis without affecting the key qualitative predictions of the model.

In our experiment, the sensory observation is an offset version of the true state, defined as:

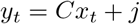

where *j* denotes the experimentally introduced jitter, i.e., the angular offset of the visual landmark only applied to the heading observation. However, in a more general case, the jitter can be applied to both heading and velocity.

In the simplest case, only heading is observed, so the observation matrix *C* is defined as

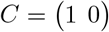

so that the jitter is applied only to the heading observation.

The predicted sensory input is

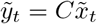

where 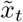 denotes the predicted state at time *t*. The resulting prediction error *δ*_*t*_ describes the mismatch between the predicted sensory input for heading and velocity, 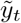, and the actual sensory input *y*_*t*_:

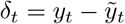

The gain matrix *L*_*t*_ describes the effect of the prediction error *δ*_*t*_ on the internal state estimates. In general, *L*_*t*_ can vary over time and as a function of the prediction error. In our task where the feedback is available only for heading, *L*_*t*_ is a vector

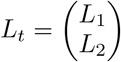

so that heading prediction error can update both the heading estimate (*L*_1_) and the velocity estimate (*L*_2_).

#### Controller: Control signal toward goal

The controller describes the control signal that drives the estimated state toward the goal. The participant’s initial estimate of the state is

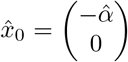

where 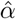 is the participant’s remembered goal angle. In general, this estimate may not be accurate (i.e., 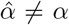), for example, due to an incorrect memory for the goal. For this simplified model, we assume no memory error, such that 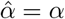.

The goal of the controller is to return to the original heading 0 and stop moving; i.e.,

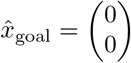

An intuitive way to reach this goal is to have a control signal that points towards this goal. In a linear model, this implies

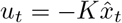

where the negative sign indicates that acceleration is directed toward the goal. Here *K* is the feedback gain

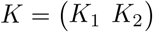

Here, *K*_1_ determines how strongly heading error contributes to the control signal, whereas *K*_2_ determines how strongly velocity error contributes to the control signal.

Substituting this control signal into the real-world dynamics gives

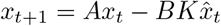

However, human turning are unlikely to be generated by an unbounded linear controller. In participant data, velocity increases during the early part of the turn, remains high during the middle of the movement, and then decreases as the participant approaches the goal (Figure 4d). This movement profile suggests that realistic motor output is bounded, such that participants cannot generate arbitrarily large angular acceleration. We therefore implement a nonlinear controller in which the control input is passed through the hyperbolic tangent function, tanh, thereby limiting the maximum acceleration:

**Fig. 4.**
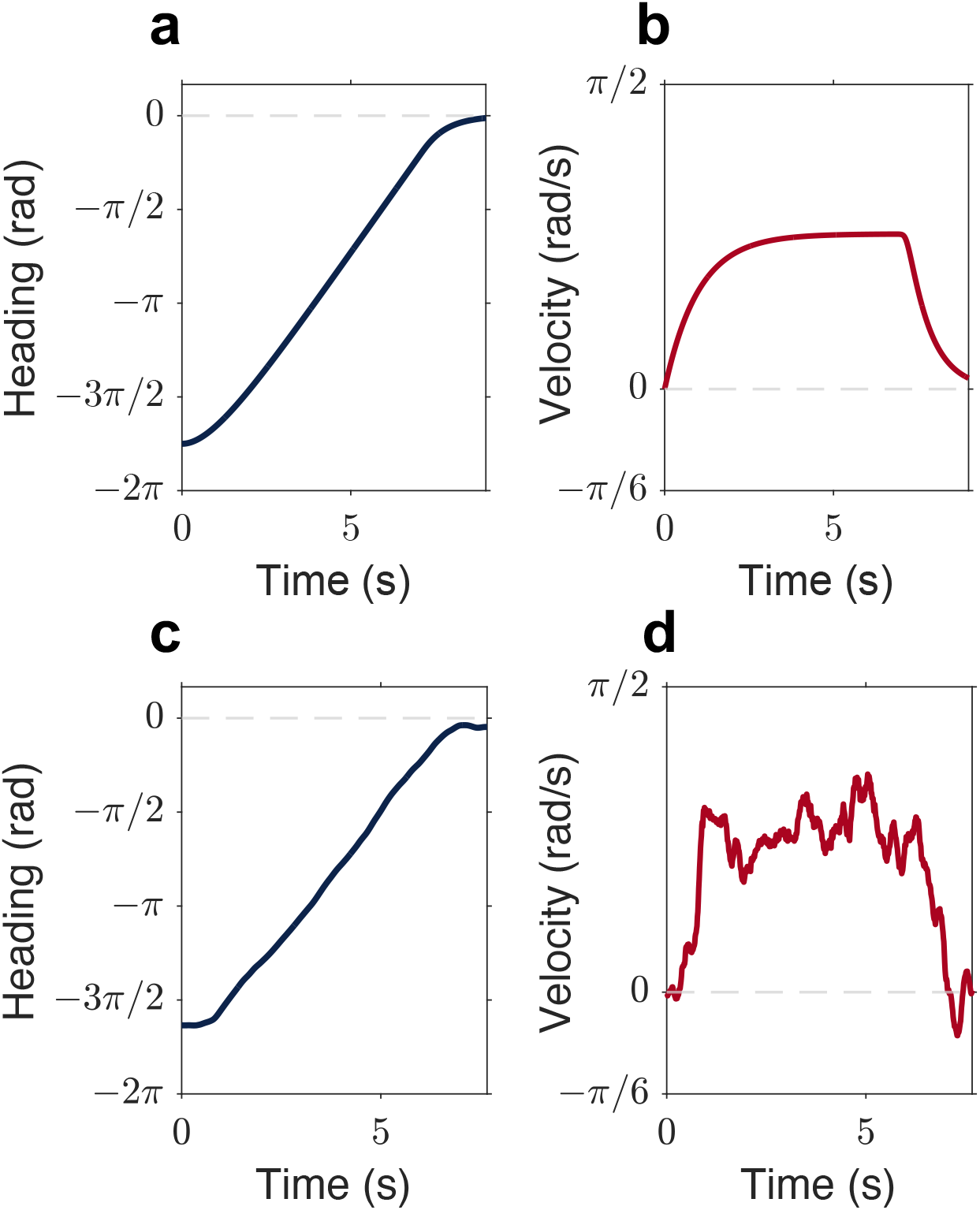
Time evolution of heading and velocity in the No Feedback condition. (a) Simulated heading trajectory. The model starts from an initial condition of −7*π/*4 and evolves smoothly toward the goal at 0. **(b) Simulated velocity profile**. Velocity increases during the early phase of the turn, remains elevated during the middle of the movement, and decreases as the system approaches the goal. **(c) Human heading trajectory**. An example trial is shown. Heading evolves smoothly toward the goal, qualitatively paralleling the simulated trajectory. **(d) Human velocity profile**. Velocity exhibits initial acceleration followed by deceleration, with a relatively high-velocity middle phase. This participant movement profile motivates a bounded-control formulation.

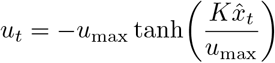

This control law is approximately linear when the desired control signal is small relative to *u*_max_, but saturates when the desired control signal becomes large. Thus, *u*_max_ represents the maximum control amplitude, or the maximum angular acceleration the participant can generate. Importantly, bounded motor output alone is not sufficient to shape the full movement profile: it constrains the magnitude of acceleration, while the resistance coefficient *β* determines the steady-state velocity at which accelerating and dissipative forces balance. Together, bounded acceleration and resistance are sufficient to produce smooth velocity profiles with a plateau, qualitatively paralleling model simulations (Figure 4b) and participant behavior (Figure 4d), without requiring an explicitly pre-planned trajectory.

This bounded-control assumption generates a testable pattern. For small reproduced angles, the required control signal is relatively small, so midpoint velocity should increase approximately linearly with reproduced angle. For larger reproduced angles, the required control signal approaches the motor limit, so midpoint velocity should plateau. To examine this pattern, we estimated midpoint velocity by averaging velocity between 25% and 75% of the movement duration, which captures the high-velocity middle portion of the turn while avoiding the initial acceleration and final deceleration phases. Midpoint velocity increased with reproduced angle for small turns but tended to saturate for larger turns (Figure 5). This plateau is consistent with bounded motor output and supports using a saturated controller in the model.

**Fig. 5.**
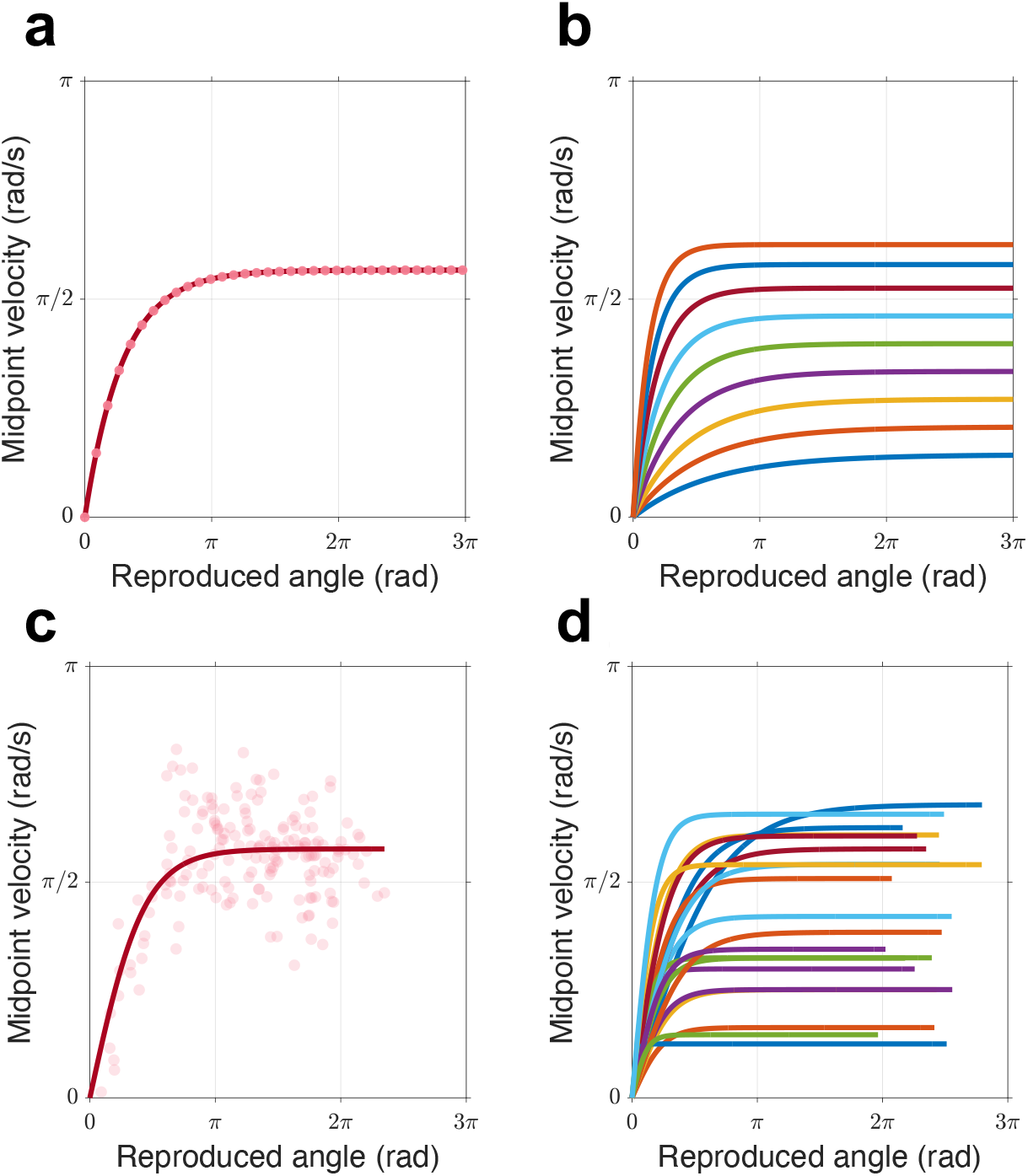
Midpoint velocity as a function of reproduced angle. **(a) Simulated midpoint velocity**. The model simulation shows that midpoint velocity increases approximately linearly for small reproduced angles but plateau for larger angles, reflecting saturation of the control input. **(b) Simulated midpoint velocity across control limits**. Varying *u*_max_ produces a family of curves with different asymptotic velocities, showing that the plateau depends on the maximum control amplitude. **(c) Midpoint velocity of one subject**. Each point represents a single trial. Midpoint velocity was estimated by averaging velocity between 25% and 75% of the movement duration. The solid line shows a fitted hyperbolic tangent function, *y* = *A* tanh(*βx*). **(d) Hyperbolic tangent fits across participants**. Despite individual differences in slope and asymptotic height, midpoint velocity tends to plateau as reproduced angle increases. This pattern is consistent with bounded motor output and motivates the nonlinear control formulation.

#### 2.2.1 No Feedback Condition: endpoint error and turning dynamics emerge from internal dynamics

In the absence of feedback, no sensory observations are available (i.e., *δ*_*t*_ = 0), and the participants rotate only based on their internal estimate of the remembered goal angle. Under the assumption that 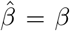, the internal estimate can be expressed as a linear transformation of the real-world state, scaled by two gain parameters: *γ*_*A*_, which captures gain on internal velocity estimation, and *γ*_*B*_, which captures gain on internal acceleration estimation. Here, *α* denotes the true target angle, and 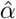 denotes the remembered angle. The full derivation is provided in Methods 4.4.1. Under this transformation, the equilibrium heading (i.e., the final reproduced angle) converges to

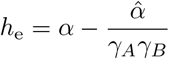

This expression shows why endpoints alone cannot identify the control process. The equilibrium heading depends on the internal gains *γ*_*A*_*γ*_*B*_, and not on the control gain *K*. Intuitively, the endpoint is determined by where the closed-loop system converges, *x*_*e*_; the controller gain determined the path to reach that equilibrium. This observation parallels the prior observer models, in which endpoint heading is determined by cue-combination parameters, such as the Kalman gain, and by the sensory perturbation [15]. In our model, the same conclusion is derived from a closed-loop dynamical system linking state estimation and motor control. This motivates modeling full movement trajectories, which contain information about the control process that is not recoverable from endpoint responses alone [30].

The dynamics of this system can be visualized in phase space (heading vs. velocity), where trajectories follow a vector field defined by the internal model and converge toward the equilibrium point *x*_*e*_. Because of gains in the internal state, this equilibrium point does not always coincide with the intended goal state *x*_goal_ (marked by a blue star), resulting in a systematic endpoint error (Figure 10b).

When plotted over time, these dynamics produce smooth velocity profiles in which the agent accelerates and then decelerates as it approaches the perceived goal (Figure 4). These movement profiles emerge naturally from the closed-loop interaction between state estimation and feedback control, without an explicit pre-planned trajectory (Figure 4).

#### 2.2.2 Feedback condition: prediction errors update state and control

When feedback is available, a prediction error *δ*_*t*_ arises from the discrepancy between the predicted and observed state at feedback onset. Let *h*_*fb*_ denote the heading at feedback onset, and let *j* denote the visual perturbation applied to the feedback signal. The prediction error updates the internal estimate with gain *L*_1_, so that the final heading reflects a weighted combination of path integration and feedback-driven correction

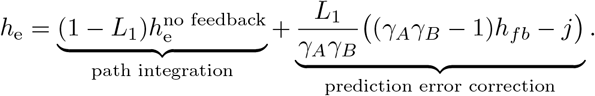

This result shows that, under feedback, the equilibrium (final) heading depends on both the feedback gain *L* and the sensory perturbation *j*, but remains independent of the control parameters *K*. Thus, similar to cue combination models, the endpoint is governed by how prediction errors update the internal state estimate, rather than by the controller itself. In particular, [15] showed that endpoint error depends on cue-combination parameters, such as the Kalman gain, and the size of the prediction error. Here, despite deriving from a closed-loop dynamical system, we arrive at the same conclusion: final position reflects the equilibrium state *x*_*e*_, which can be shifted by internal estimation biases and feedback-driven corrections. See full derivation in Methods 4.4.1.

Mostly importantly, the control model introduces a time-resolved prediction. By integrating prediction errors into the state estimate, the model can generate systematic overshoot or undershoot and time-resolved changes in control signals during movement. Specifically, the model simulation predicts that prediction error should induce an immediate change in the control signal, manifesting as a change in acceleration following feedback onset (Figure 6). These trajectory-level predictions distinguish the proposed model from the previous endpoint observer models, which explain final position but do not explicitly model when and how prediction errors update the control dynamics that generate the movement trajectory.

**Fig. 6.**
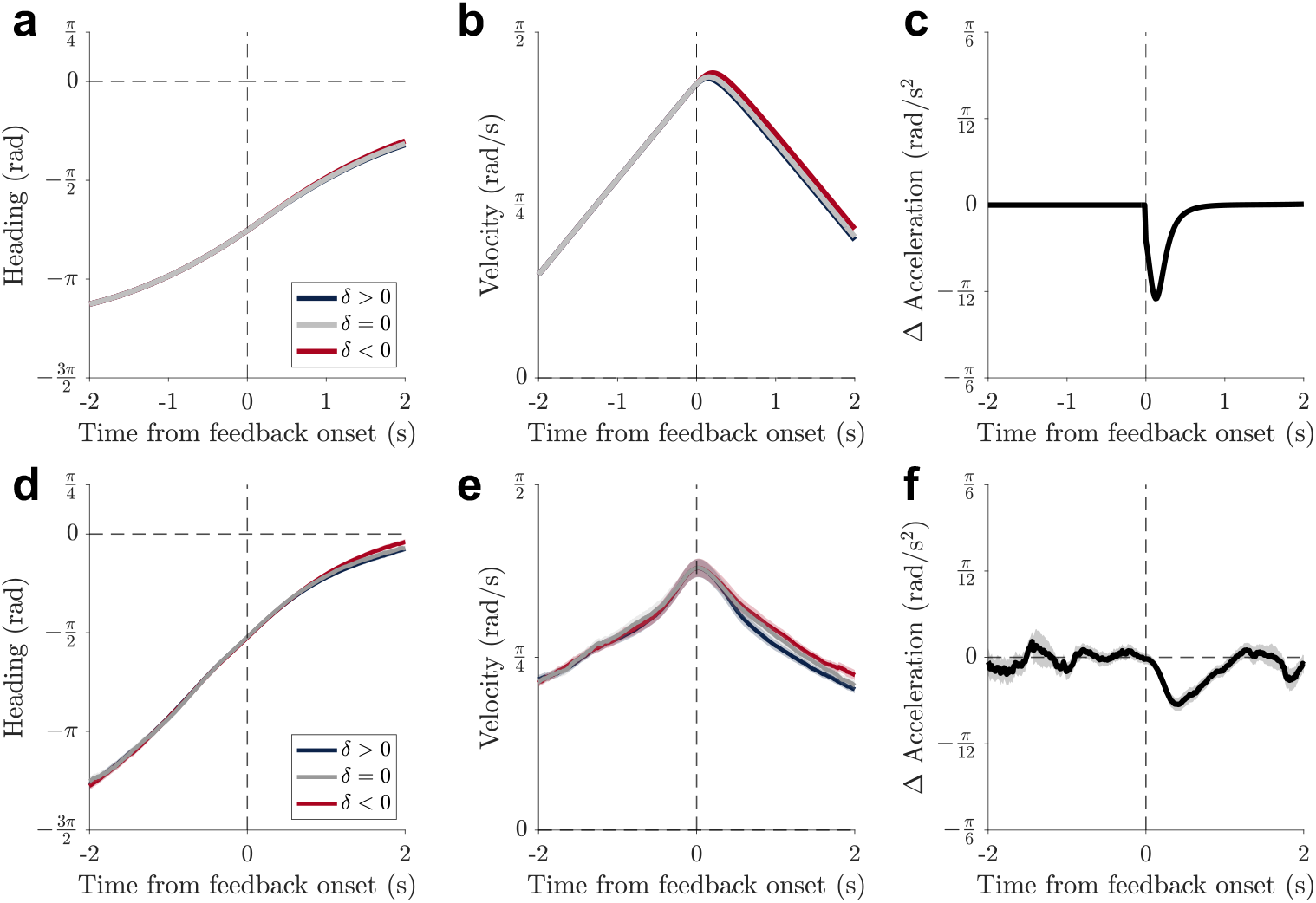
Prediction errors induce changes in model simulations and participant behavior. (a) Model heading. Positive (*δ >* 0), zero (*δ* = 0), and negative (*δ <* 0) prediction errors produce different simulated heading trajectories following feedback onset (*t*_fb_). **(b) Model velocity**. Prediction errors modulate the timing and magnitude of simulated velocity profiles. **(c) Model acceleration**. Simulated acceleration diverges as a function of prediction error. The black line shows the divergence between positive and negative conditions (Δ*a* = *a*_*δ>*0_ − *a*_*δ<*0_), highlighting the predicted control signal. **(d) participant heading**. Average heading trajectories after feedback onset show divergence across prediction error conditions. Shaded regions represent *±*SEM. **(e) participant velocity**. Velocity profiles exhibit prediction-error–dependent modulation consistent with the model. *±*Shaded regions represent *±*SEM. **(f) participant acceleration**. Acceleration shows a time-locked divergence following feedback onset. The black line (Δ*a*) reflects the acceleration difference associated with prediction error, qualitatively paralleling the model simulation. Shaded regions represent *±* SEM.

### 2.3 Prediction errors induce time-locked acceleration changes consistent with closed-loop control

A key prediction of the model is that discrepancies between predicted and observed states generate prediction errors that update the control signal. Unlike observer models such as [15], this model makes a specific prediction in the time domain: visual prediction errors should induce an immediate correction in the control signal, manifesting as a change in acceleration following feedback onset, as shown in Figure 6. The model simulations shown in Figure 6a–c were generated using the simulation parameters reported in Table 1; they are not fitted trajectories.

**Table 1.** Simulation parameters for Figure 6a–c.

| Parameter | Value | Description |
| --- | --- | --- |
| $dt$ | 0.01 s | Simulation time step |
| $N$ | 400 | Number of simulation steps |
| $t_f$ | 200 | Feedback onset step |
| $u_{\max}$ | 0.67 | Maximum control input |
| $x_0$ | 3.7 rad | Initial heading magnitude |
| $\gamma_A$ | 1.2 | Velocity gain |
| $\gamma_B$ | 1.0 | Acceleration gain |
| $K_1$ | 3.8 | Heading feedback gain |
| $K_2$ | 5.4 | Velocity feedback gain |
| $\beta$ | 0.001 | True resistance coefficient |
| $\hat{\beta}$ | 0.001 | Estimated resistance coefficient |
| $L_1$ | 0.09 | Heading Kalman gain |
| $L_2$ | 0 | Velocity Kalman gain |
| $\delta$ | $\pm 1.3$ rad | Prediction error |

Simulations show that the closed-loop model is sufficient to generate systematic changes in heading, velocity, and acceleration trajectories after prediction error (Figure 6a–c). In particular, the model produces a divergence in acceleration between positive and negative prediction-error conditions, providing a qualitative signature of control adjustment.

We next examined whether similar patterns were observed in participant behavior. Movement dynamics around feedback onset showed qualitatively similar patterns. Heading and velocity trajectories differed across prediction-error conditions (Figure 6d–e). More importantly, acceleration exhibited a time-locked divergence following feedback onset (Figure 6f). Before feedback, the acceleration difference between positive and negative prediction-error trials was small (mean difference = 0.009 rad/s^2^). After feedback, the difference increased in magnitude (mean difference = −0.117 rad/s^2^). To quantify this divergence, we fit a linear mixed-effects model with prediction-error sign, time window, and their interaction as fixed effects and participant as a random intercept. The analysis included 172 sessions from 107 unique participants; four sessions were excluded because acceleration estimates were not available due to short trial duration. The Prediction Error Sign × Time Window interaction was significant (*β* = −0.126 rad/s^2^, SE = 0.064, 95% CI [− 0.252, 0.00003], *z* = −1.96, *p* = 0.050), indicating that acceleration profiles for positive and negative prediction-error trials diverged following visual feedback.

This pattern is consistent with the model prediction that prediction errors are associated with changes in control signals. The result should not be interpreted as a quantitative validation of the model, but it suggests that rotational navigation behavior reflects a dynamical process in which sensory mismatches are linked to adjustments in ongoing movement.

## 3 Discussion

Our results show that visual prediction errors are associated with time-locked changes in angular acceleration during turning, consistent with a control-theoretic model in which internal state estimates are updated to guide motor output. These findings address a gap in the navigation literature by linking heading estimation with continuous motor behavior. Specifically, the proposed model bridges two lines of research that have largely developed in isolation: observer models of cue combination in navigation [12–18, 31] and controller models of motor control [23–26, 28].

A major advantage of this framework is that it incorporates existing observer models as a special case while extending them to explain movement dynamics (see Results 2.2.1). Prior work has established that visual landmarks affect final stopping position during path integration [14, 15, 18, 31]. Our endpoint results reproduce this finding, confirming that participants incorporated visual landmarks into their final responses (see Fig 2). In this sense, the current model preserves the key contribution of existing observer models by capturing how visual and self-motion cues are combined to influence final responses.

Importantly, the closed-loop model goes beyond endpoint behavior by showing how motor commands can affect the trajectory used to reach the goal. Prior observer models account for where participants stop, but do not specify the control dynamics of the journey and therefore cannot predict how velocity and acceleration evolve during movement. Our simulations show that explicitly modeling heading and angular velocity as jointly estimated states can account for the dynamics of movement during angle reproduction. The proposed model therefore generates not only predictions about the final heading, but also predictions about the time evolution of heading and velocity during turning. The simulation and participant trajectories show smooth heading changes and velocity profiles with initial acceleration, a high-velocity middle phase, and final deceleration (see Fig 4). Movement trajectories thus reveal dynamic control structure that is obscured when continuous trajectories are reduced into a single endpoint [30]. Consistent with this idea, recent work in naturalistic walking tasks has similarly shown that temporal dynamics carry information beyond what endpoints alone provide [19, 27, 32].

A second contribution of the framework is that it explicitly models the control signals generated during movement. Prior observer models explain how sensory cues update internal estimates, but do not specify how these updated estimates are transformed into motor output over time. In contrast, the proposed model shows how prediction errors can update the estimated state, which in turn changes the control signal during ongoing movement. This qualitative prediction is consistent with human data: visual feedback was followed by time-locked changes in angular acceleration (see Fig 6), suggesting that landmark mismatches are linked to online motor corrections. As a closed-loop model, the framework also provides a way to ask how multisensory inputs are translated into control signals during ongoing behavior, rather than only whether participants rely on a given sensory cue.

Three limitations should be noted. First, the model is qualitative in the manuscript and thus should be interpreted as an existence proof. Several parameter combinations can produce similar model behaviors, and the current analyses do not estimate participant-wise parameters for estimation gains, feedback gains, or motor limits. The value of the model is therefore to serve as a testable hypothesis where a closed-loop architecture separating state estimation from feedback control is sufficient to generate qualitative dynamics observed in participant turning, including endpoint biases, bounded velocity profiles, and feedback-locked acceleration corrections. This approach is useful because it makes explicit which future measurements would distinguish estimation errors from control errors. Second, the current paradigm uses brief visual feedback rather than continuous sensory feedback. This design isolates the effect of prediction error from landmarks, but it may not fully capture heading estimation in richer environments where visual, vestibular, proprioceptive, and motor signals are continuously available [5]. Third, although the observed behavioral dynamics are consistent with a closed-loop control interpretation, the study does not directly measure the neural mechanisms that implement these computations. Therefore, any mapping between prediction-error-driven control signals and neural circuits remains a hypothesis to be tested in future work.

This control-theoretic framing situates the present work within a broader effort in computational neuroscience and cognitive science to model behavior as a continuous, dynamic, and coupled perception–action process over time. Feedback-control accounts of biological behavior emphasize that sensing and action are not independent stages, but mutually coupled processes embedded in ongoing interaction with the environment [33]. Neurocybernetic approaches similarly argue for interpretable dynamical models that link brain, body, and environment, especially when behavior unfolds through latent states that cannot be inferred from endpoints alone [34]. In navigation, probabilistic models have shown that human navigation strategies and errors can emerge from dynamic interactions among spatial uncertainties, self-motion estimates, landmark information, planning, and action execution [19]. Bayesian accounts of ring attractor networks further show how directional representations can implement uncertainty-sensitive state estimation over time [16]. The present model builds on these ideas by focusing on rotational navigation as a closed-loop observer–controller problem: visual landmark mismatch updates an internal heading estimate, and the updated estimate is transformed into evolving velocity and acceleration through feedback control. Thus, the contribution of the present study is to connect endpoint-level observer models of heading estimation with trajectory-level models of motor control, showing that continuous movement dynamics can reveal control dynamics that are obscured when behavior is measured only by where participants stop.

## 4 Methods

### 4.1 Participants

A total of 105 paid undergraduate students (age range 18–21 years) from the University of Arizona participated. Data collection occurred in two phases: an initial cohort of 34 students (17 female, 17 male) and a second cohort of 71 students (40 female, 31 male). Participants in the second cohort each completed two sessions, resulting in a total of 176 sessions across the combined dataset. Participants in the first cohort were compensated using funds from the University of Arizona Research and Project (ReaP) Grant. Participants in the second cohort were compensated using funds from the SensorLab Seed Grant. All participants provided informed consent prior to participation.

### 4.2 Apparatus and Stimuli

The experimental task was developed in Unity (version 2018.4.11f1) using the Landmarks 2.0 framework [35]. Participants wore an HTC Vive Pro head-mounted display equipped with a wireless adapter and held a pair of HTC Vive controllers (Figure 1a). Participants were positioned at the center of a rectangular virtual room (13 m × 10 m × 5 m). The environment was designed to appear naturalistic and contained multiple stable visual landmarks to support heading judgments (Figure 1a). Head position and velocity were recorded at 90 Hz for each trial. The use of immersive virtual reality enables controlled yet naturalistic investigation of participant navigation, allowing researchers to manipulate sensory inputs while preserving active, embodied movement that engages proprioceptive and vestibular signals [36].

### 4.3 Rotation Task

Each participant completed 200 trials organized into three blocks: 10 no-feedback trials, 180 feedback trials, and 10 no-feedback trials (Figure 1a). During the encoding phase, participants were guided to rotate to an angle ±*α* using haptic cues delivered through the handheld controllers. Vibration of the left controller cued leftward rotations, and vibration of the right controller cued rightward rotations. The vibration stopped once the encoded angle was reached. During encoding, full visual feedback was available, allowing participants to integrate visual, vestibular, and proprioceptive signals to form an internal estimate of angular displacement.

During the reproduction phase, participants were instructed to rotate back toward the starting heading 0 by rotating in the opposite direction. In the No-Feedback condition, no visual feedback was available during reproduction (Figure 1b). The reproduced angle reflected the participant’s internal estimate of angular displacement, denoted as 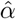, derived from path integration. In the Feedback condition, participants received brief visual feedback (300 ms) at time *t*_*fb*_ (Figure 1c). This feedback could be either consistent or inconsistent with the participant’s true heading at that time, introducing a mismatch between predicted and observed heading (visual prediction error). A positive *j* corresponded to the virtual room appearing further ahead in the direction of movement than the participant’s true heading, thereby producing a positive prediction error. For example, in Fig. 1b, although the participant had not yet aligned with the center of the right wall, the visual scene suggests that they had already reached it. Thus, the participant reduced further turning relative to the no-feedback condition. This manipulation dissociated self-motion cues from visual landmarks, allowing us to examine how visual prediction error influenced the updating of internal heading estimates. Participants rotated at their own pace without haptic guidance. When they believed they had returned to the initial heading 0, they pressed a controller button to register their response, and the final response angle was recorded. The response error was defined as the difference between the response angle and the encoded angle.

The key experimental manipulation in this study was whether visual feedback was presented during the reproduction phase and, if so, how informative it was. In the No-Feedback condition, participants viewed a blank screen during reproduction and relied only on path integration. In the Feedback condition, participants briefly viewed the room for 300 ms at time *t*_*fb*_, with a jitter angle *fb* that was either consistent or inconsistent with their current heading. Consistent feedback occurred with probability *ρ* = 0.7. In this case, the jitter angle was sampled from a Gaussian distribution centered on the true heading with a standard deviation of 30°. Inconsistent feedback occurred with probability 1 − *ρ* = 0.3. For the 34 participants in the one-session dataset, inconsistent feedback was sampled from a uniform distribution between −100° and +100° (− 5*π/*9 to + 5*π/*9 rad). For the 71 participants in the two-session dataset, inconsistent feedback was sampled from a wider range, −180° to +180° (− *π* to + *π* rad). Because the smaller jitter range made the visual feedback less extreme, participants in the one-session cohort were less likely to ignore the visual feedback entirely, whereas the wider range in the two-session cohort allowed some feedback events to be sufficiently misleading that participants could down-weight or ignore them. Because some participants rotated more than one full circle during reproduction, all angular variables were analyzed in radians to preserve continuity at the 0°*/*360° boundary.

Formally, the jitter angle *f* was sampled according to:

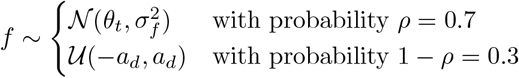

where *a*_*d*_ = 100° = 5*π/*9 for the one-session dataset and *a*_*d*_ = 180° = *π* for the two-session dataset.

This feedback design ensured that visual information was sufficiently informative to encourage reliance on it, yet variable enough to probe the impact of misleading feedback across the full angular space. To further encourage participants to use the visual feedback, they were not informed that the feedback could be misleading.

### 4.4 Control Theory Model of Heading

Our goal in this section is to build a qualitative control-theoretic model of heading over time as people turn back to the goal. The model follows the logic of linear-quadratic-Gaussian and related feedback-control frameworks by separating the observer, which estimates the current state from internal dynamics and sensory prediction error, from the controller, which converts the estimated state into a motor command. This separation is useful even without quantitative fitting because it makes explicit how sensory uncertainty, prediction error, and motor correction can interact over time. We begin with a noiseless discrete-time linear controller with no time delays. The linear model gives insight into the behavior but leads to unrealistically large turning speeds in some conditions. This high-velocity flaw can be addressed by adding a saturating nonlinearity to the control signal, which we do in the next subsection. Because the current dataset is used for qualitative model assessment rather than parameter identification, the simulations should be interpreted as an existence proof that this constrained architecture can generate participant-like movement dynamics. All analyses and simulations were performed in MATLAB R2024b.

#### 4.4.1 Linear Models

##### Real-world state

The real-world state describes the physical dynamics of turning. For simplicity, we quantify turning behavior via two variables: the heading angle *h*_*t*_ and angular velocity *v*_*t*_. In the Rotation Task, participants generate a control input *u*_*t*_, which manifests as an angular acceleration. The continuous system can be approximated in discrete time via Taylor expansion:

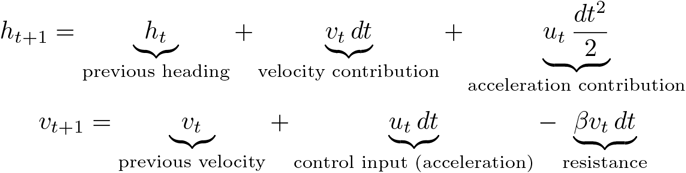

where *dt* is the time step and *β* ≥ 0 is a resistance coefficient. These equations reflect that heading evolves as the time integral of angular velocity, while angular velocity is driven by the control input and opposed by velocity-proportional resistance.

Because the contribution of acceleration to position over a single time step (*dt* = 0.01, corresponding to the sampling interval of the HTC system) is small, we neglect the second-order term *u*_*t*_ *dt*^2^*/*2 in the heading update. This yields

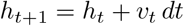

We now express these dynamics in state-space form. Turning behavior is quantified by two variables: the heading angle *h*_*t*_ and its velocity *v*_*t*_, which correspond to the data measured in the experiment. Together, they form the state vector

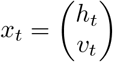

In the Rotation Task, the participant’s goal is to move this state vector from the encoded angle −*α* to the door at angle 0, in both locations standing still so that the angular velocity is 0 (Figure 1).

Without loss of generality, we define the door as the goal state

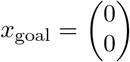

which corresponds to zero heading error and zero angular velocity. Here *x*_goal_ denotes the intended controller goal, whereas *x*_*e*_ denotes the equilibrium of the closedloop system. All state variables are expressed relative to the intended goal. Specifically, any goal can be shifted to the origin via a change of variables 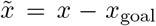 [37]. Under this transformation, the system dynamics are preserved, and the control problem becomes one of stabilizing the system to the origin, i.e., driving both heading and velocity to zero. The observed endpoint corresponds to *x*_*e*_, which may differ from *x*_goal_ when the internal estimate is biased or when feedback changes the estimated state.

Under this transformation, the initial state becomes

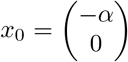

To reach the goal, participants generate a control signal *u*_*t*_, which updates the state according to

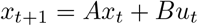

The matrices *A* and *B* follow directly from the discretized physical dynamics above. In particular, the state transition matrix *A* is written as

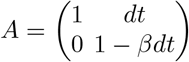

which reflects that heading integrates velocity, and velocity decays over time due to resistance. And the control input matrix *B* is

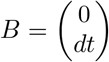

which updates angular velocity.

##### Estimated state

The estimated state describes participant’s internal estimation of the real-world states. Participants do not observe the true state of the world *x*_*t*_ directly. Instead, they must infer their heading and angular velocity by integrating self-motion cues over time and combining these internal estimates with visual and other sensory cues when they are available. We write this estimated state as

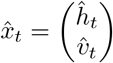

which evolves over time according to

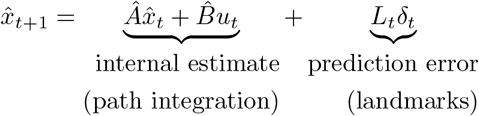

where *Â* is the participant’s internal estimate of the dynamics matrix, 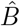 is the participant’s estimate of the effect of the control signal. *L*_*t*_ is the gain applied to sensory prediction error, and *δ*_*t*_ is the mismatch between predicted and observed sensory input. In general, *Â* ≠ *A* and 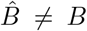, which captures the possibility that participants’ internal model of the dynamics differs from the real-world dynamics [28].

We define these matrices as

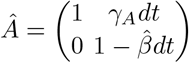

and

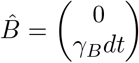

These forms assume that the participant applies gains to their internal estimates of velocity and acceleration, scaling velocity by a factor of *γ*_*A*_ and acceleration by a factor of *γ*_*B*_. Note that here *γ*_*A*_ plays a similar role as *γ*_*d*_ in [15]. Such gain parameters are consistent with prior work showing that path integration errors scale with travelled distance, reflecting systematic accumulation of error in internal motion estimates [8, 29]. In addition, for simplicity we assume that the participant’s estimate of resistance matches the true resistance, that is 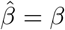. While this assumption could be relaxed, it simplifies the analysis without affecting the key qualitative predictions of the model.

In our experiment, the sensory observation is an offset version of the true state, modeled as:

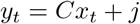

In the simplest version of the model, the observation matrix *C* is defined as the row matrix:

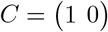

so that only the heading (position) is observed. The scalar *j* represents the experimentally introduced jitter, i.e., the angular offset of the visual landmark only applied to the heading observation. However, in a more general case, the jitter can be applied to both heading and velocity.

The prediction is

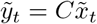

where 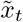 is the internal model’s prediction of the state at *t*.

The prediction error *δ*_*t*_ describes the mismatch between the predicted sensory input for heading and velocity, 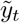, and the actual sensory input *y*_*t*_:

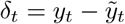

The Kalman gain matrix *L*_*t*_ describes the effect of the prediction error *δ*_*t*_ on the internal state estimates. In general, *L*_*t*_ can vary over time and as a function of the prediction error. In the case where the feedback is only position (and not velocity), *L*_*t*_ is a vector

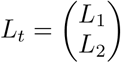

describing the effect of the position prediction error on the heading (*L*_1_) and the velocity (*L*_2_).

##### The control signal

The controller describes the control signal that drives the system to the goal based on the internal estimate of state and goal. The initial condition for the participant’s estimate of the state is

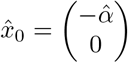

where 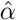 is the participant’s estimate of the goal. In general, this estimate may not be accurate (i.e., 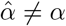) due to errors induced, for example, by an incorrect memory for the goal. For this simplified model, we will assume that there are no memory errors such that 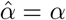.

The goal of the controller is to get back to facing the door and stop moving; i.e.,

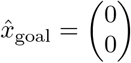

An intuitive way to get to this goal is to have a control signal that points towards this goal. In the linear case, this implies participant generates control inputs based on the estimated state:

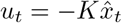

where the negative sign points the acceleration towards the goal and *K* is a 1 × 2 feedback gain vector

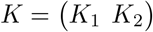

where *K*_1_ describes the effect of position mismatch on the control signal and *K*_2_ describes the effect of the velocity mismatch.

Substituting the expression for the control signal into the update equations for the dynamics we get, for the real-world dynamics,

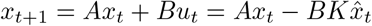

##### Modeling the task in the no-feedback condition

In the no feedback condition, when 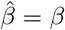, there exists a linear mapping from the real-world state *x*_*t*_ to the internal state estimate 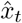. This mapping reduces the dynamics from the four-dimensional space spanning *x*_*t*_ and 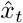 to the two-dimensional space spanning *x*_*t*_.

To compute this mapping, we note that the only distortions in the internal model are the initial condition and the scalings on the velocity and acceleration.

Thus, the internal estimate of the velocity is simply a scaled version of the real-world velocity

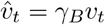

Likewise, the distance traveled in the internal model, 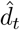, is simply a scaled version of the distance traveled in the real world, *d*_*t*_

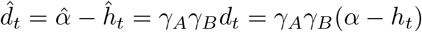

*Note:* These relationships hold regardless of the control signal, so long as the subject’s estimate of the resistance is the same as the true resistance (i.e., 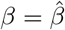).

This allows us to write down a linear relationship between the internal state 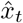 and the real-world state *x*_*t*_:

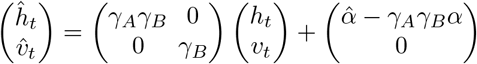

Equivalently,

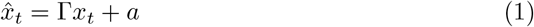

where

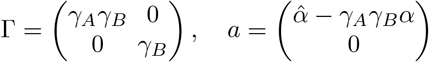

We refer to Equation 1 as the “no feedback transformation”.

If we substitute this expression into the control equation for *x*_*t*_ we get

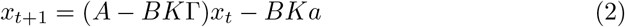

Combined, Equations 1 and 2 reduce the four-dimensional state equation down to two dimensions and we can do everything in the space of the experimentally observed state *x*_*t*_. We now consider the two-dimensional dynamics implied by Equation 2 in more detail.

##### Phase space of the dynamics

One way to understand the dynamics of Equation 2 is in a phase space—plotting heading against velocity to visualize a trajectory. An example is shown in Figure 7. In this plot, each point in phase space is associated with a vector indicating the change in heading and velocity at that point. Expressions for these vectors can be obtained by looking separately at the evolution of heading and velocity implied by Equation 2.

**Fig. 7.**
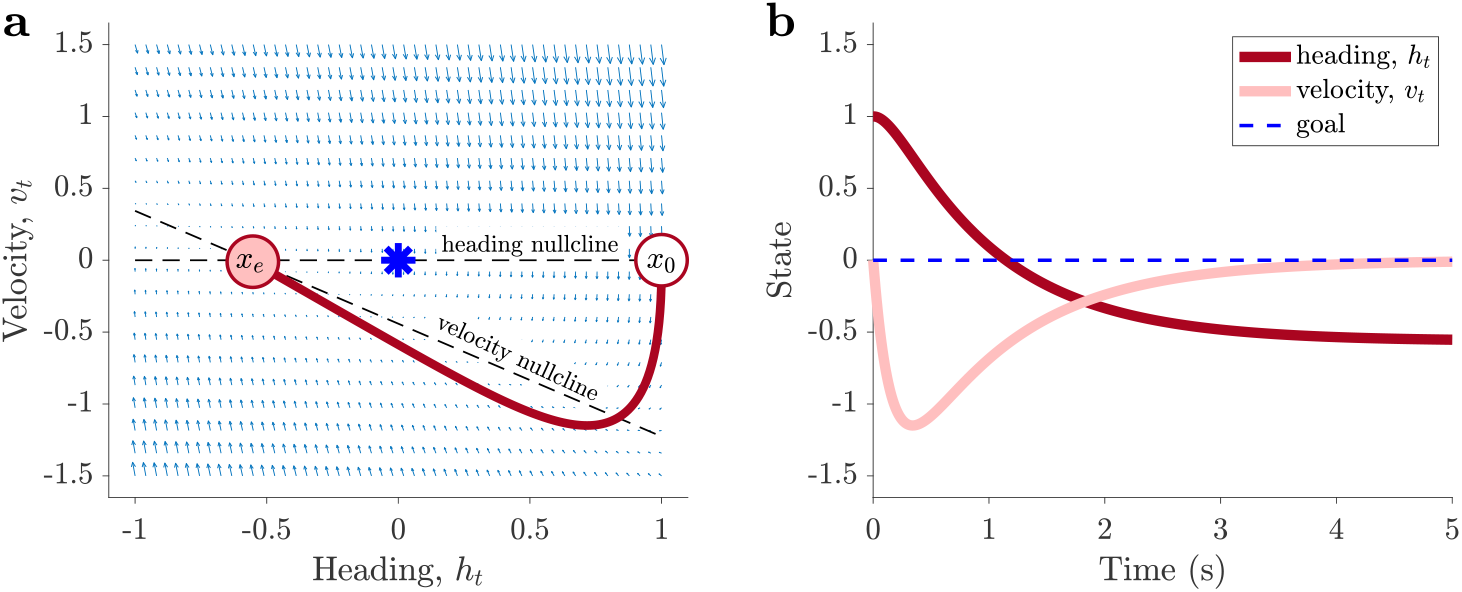
Trajectory in phase space and over time. Example trajectory in phase space (a) and over time (b) for parameters *γ*_*A*_ = *γ*_*B*_ = 0.5, *K*_1_ = 10, *K*_2_ = 8.94, *β* = 1, and *dt* = 0.01. **a)** The system evolves in a two-dimensional state space defined by heading (*h*_*t*_) and angular velocity (*v*_*t*_). Trajectories converge toward an equilibrium point *x*_*e*_ determined by the intersection of nullclines. In this example, errors in path integration cause *x*_*e*_ to differ from the intended goal state *x*_goal_ (blue star), resulting in an endpoint heading error. **b)** The same trajectory plotted as a function of time.

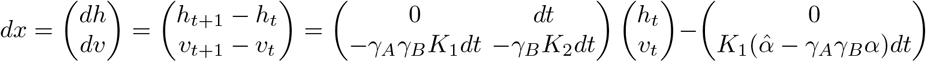

So

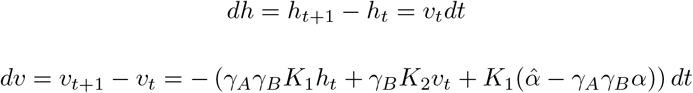

This implies the nullclines, defined as *dh* = 0 or *dv* = 0 in phase space (Figure 7). The intersection of these two nullclines defines *x*_*e*_, the equilibrium point of the system.

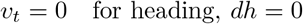

and

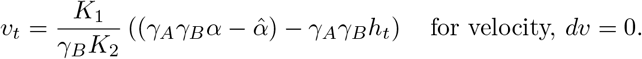

##### Solution for state as a function of time

If we write

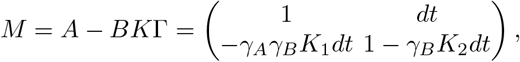

we can solve this equation to compute the state at any point in time

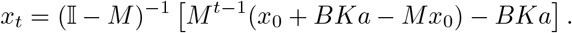

Or, if we make the identifications that the state at *t* = 1 is

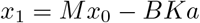

and the equilibrium state is

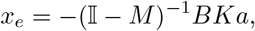

then we have

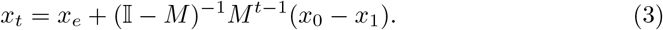

##### Final heading does not depend on K

Because *M* is a 2×2 matrix, computing the inverse in this equation is straightforward:

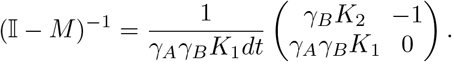

and we can write down the equilibrium state as

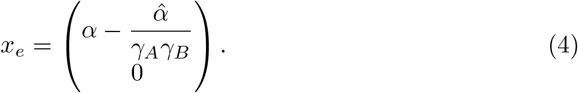

Thus, the equilibrium heading that this model turns to only depends on *γ*_*A*_*γ*_*B*_ and not on the control parameters *K*_1_ and *K*_2_.

##### Aside

There is a much easier way to obtain this result from the no-feedback transformation:

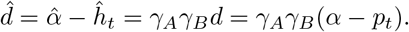

At the equilibrium, the estimated heading has reached the intended controller goal, *ĥ*_goal_ = 0, which implies

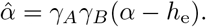

Rearranging gives

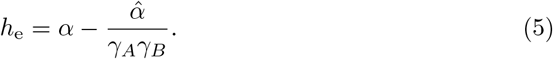

This expression may also explain [15] as a special case. There is the strong relationship observed between gain on goal and gain on velocity in [15]. If *γ*_*A*_ = *γ*_*B*_ = *γ* then the dependence becomes quadratic (*γ*^2^), which is numerically similar to a linear relationship over a limited parameter range.

##### Dynamics depends on all parameters

While Equation 3 solves the dynamics, we can gain more insight into the qualitative properties of the trajectories by computing the eigenvalues of *M* :

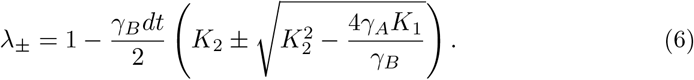

This shows that the eigenvalues, and hence the dynamics, depend on all parameters in the model.

Moreover, the eigenvalues will be complex when

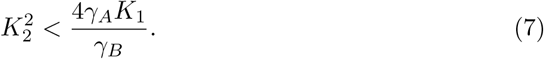

In this regime, the dynamics exhibit oscillations, implying overshoot in the trajectory — i.e., the agent will turn beyond its goal before correcting.

##### Scaling law of the dynamics

Equation 3 can be rewritten to reveal a scaling law for the dynamics:

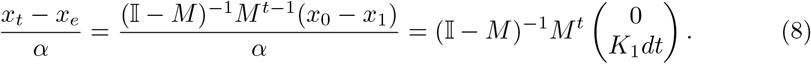

For given parameters, this shows that the quantity

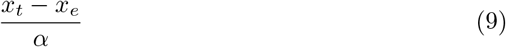

evolves identically for all starting points. This is illustrated numerically in Figure 8.

**Fig. 8.**
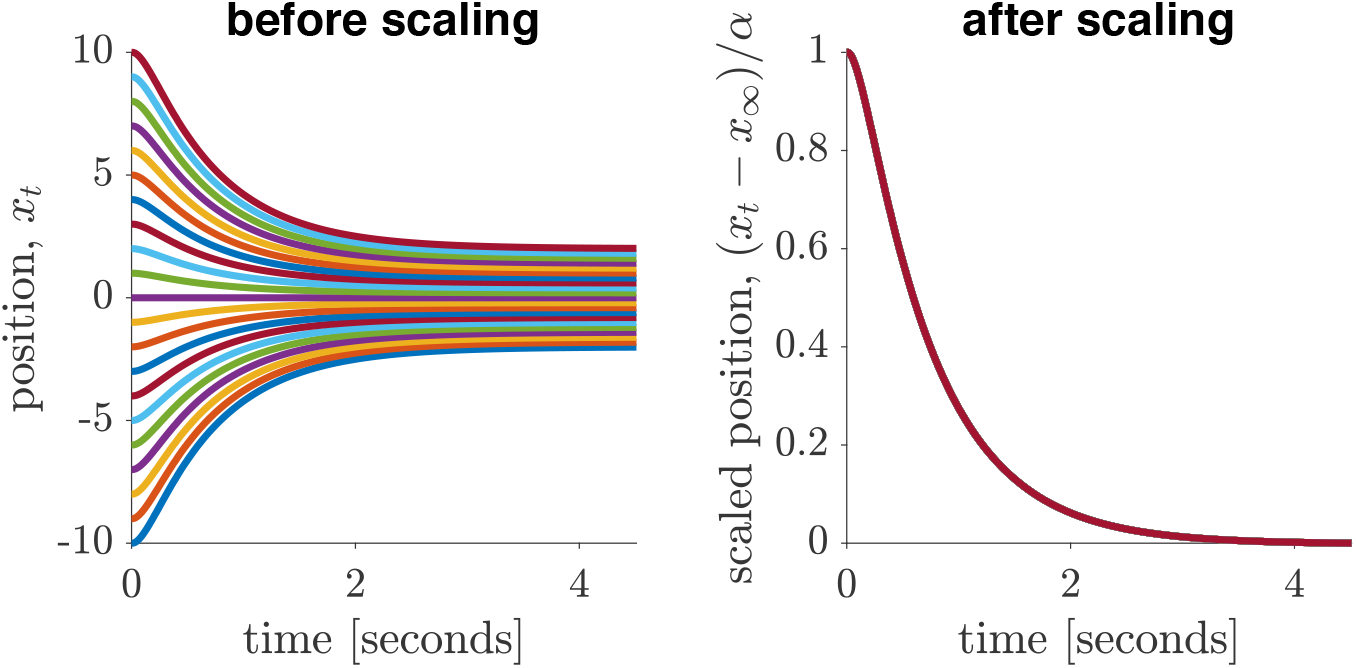
Scaling law of the dynamics. **Left:** Position over time for different starting points. **Right:** After scaling by *α*, all trajectories collapse onto a standard form.

**Fig. 9.**
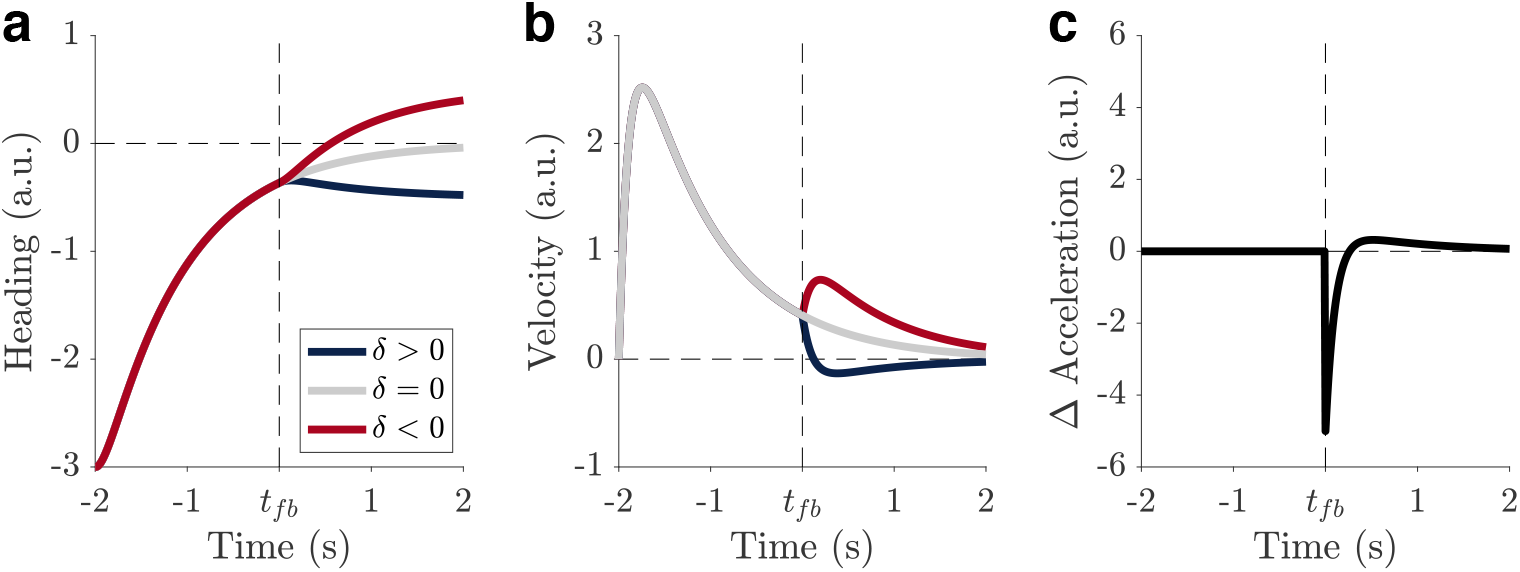
Linear model predictions for the effect of prediction error on heading, velocity, and acceleration. (a) **Heading**. Positive, zero, and negative prediction errors produce different heading trajectories following feedback onset (*t*_fb_). Blue denotes positive prediction error (*δ >* 0), grey denotes no prediction error (*δ* = 0), and red denotes negative prediction error (*δ <* 0). A positive prediction error (observation ahead of the internal estimate) means the agent believes they have traveled farther than they actually have, resulting in a final heading below the goal. In contrast, a negative prediction error (observation behind the estimate) leads to overshooting the goal. (b) **Velocity**. Velocity follows the same color convention (blue: *δ >* 0, grey: *δ* = 0, red: *δ <* 0). (c) **Acceleration**. The black line shows the difference in acceleration between positive and negative prediction error conditions (Δ*a* = *a*_*δ>*0_ − *a*_*δ<*0_), highlighting the prediction-error–dependent control signal.

**Fig. 10.**
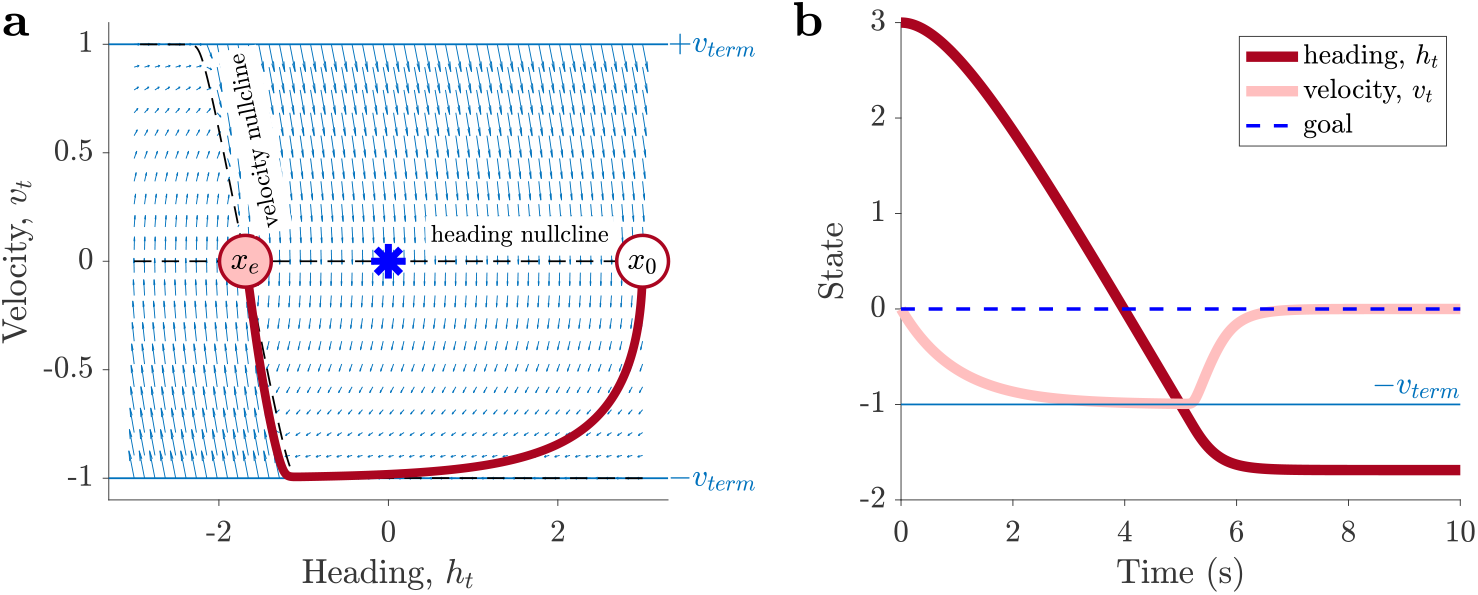
(a) Phase portrait of the nonlinear system. The system evolves in a two-dimensional state space defined by heading (*h*_*t*_) and angular velocity (*v*_*t*_). Trajectories converge toward an equilibrium point *x*_*e*_ determined by the intersection of nullclines. The curvature reflects bounded control due to a saturating tanh nonlinearity. (b) Time evolution of heading *h*_*t*_ and velocity *v*_*t*_.

This scaling law highlights one of the key limitations of the linear model. When the starting point is increased, both heading and velocity scale proportionally. Thus, there is no intrinsic bound on turning speed or acceleration in this formulation.

##### Modeling the task in the feedback condition

We model the flash of visual feedback by assuming that at one time point, *t*_*fb*_, the agent is presented with a prediction error *δ*_*t*_, which it uses to update its state estimate with gain matrix *L*. At all other times, there is no visual feedback and no prediction error.

To understand the dynamics in the case of a prediction error, we first focus on the estimated state. Before the feedback, this evolves according to

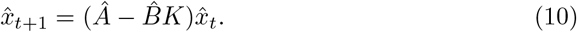

Thus, at time *t*_*fb*_, before the prediction error is applied, the estimated state is

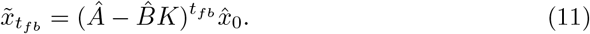

This leads to a prediction error

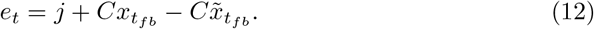

The real-world state at the time of feedback is

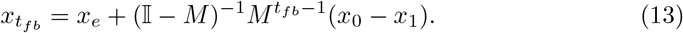

Applying this prediction error to update the estimate gives

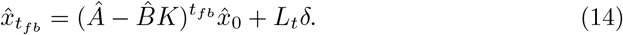

After feedback, it continues to evolve as

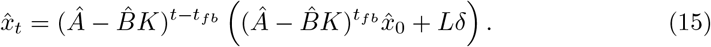

Simplifying,

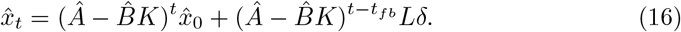

##### Solution for final heading

From the no-feedback transformation, at the time of feedback the predicted heading is

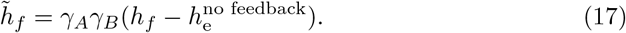

This prediction is compared with feedback *f*, creating a prediction error that updates the heading to

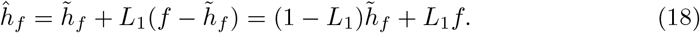

By the no-feedback transformation, the new heading obeys

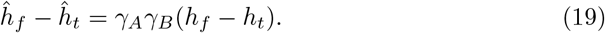

Rearranging,

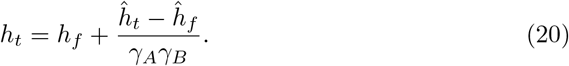

Substituting for *ĥ*_*f*_,

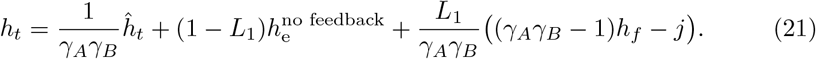

where

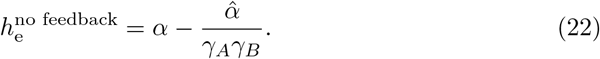

Setting *ĥ*_*t*_ = 0 gives the equilibrium (final) heading under feedback:

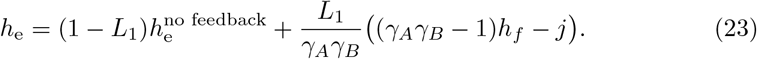

Thus, the final heading does not depend on the control parameters *K*, but does depend on the jitter and the feedback gain.

##### Control signal after feedback

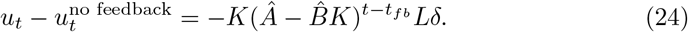

#### 4.4.2 Non-linear Model

The linear model permits unbounded velocities and accelerations. A more realistic model assumes a maximum acceleration *u*_max_. We implement this using a saturating hyperbolic tangent (tanh) nonlinearity:

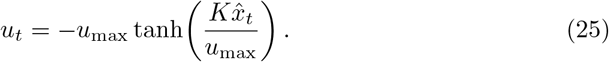

Thus, for small control input, the nonlinear model reduces to the linear model because

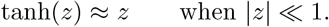

The nonlinear model differs primarily when the control input is large, in which case the acceleration saturates at ±*u*_max_.

The nonlinearity is approximately linear near zero but saturates at ±*u*_max_ for large inputs. With resistance, this leads to a terminal velocity

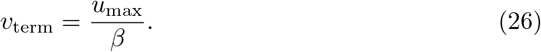

Substituting into the real-world dynamics,

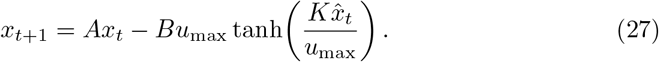

For the estimated state,

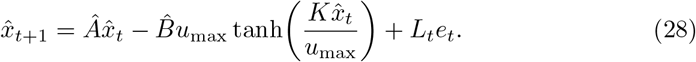

##### No feedback condition

With 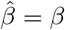, the no-feedback transformation still holds:

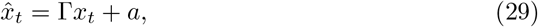

where

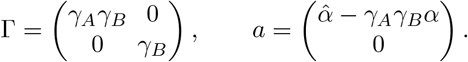

Substituting this transformation into the nonlinear control law gives a reduced two-dimensional nonlinear system in the real-world state:

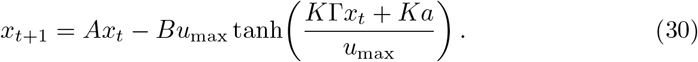

Because

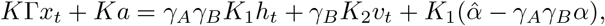

and because the no-feedback equilibrium is

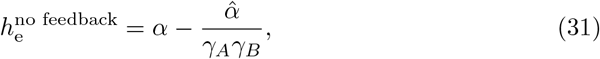

we can rewrite the control argument as

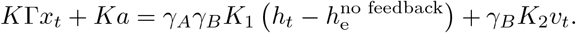

Thus, the nonlinear no-feedback dynamics are

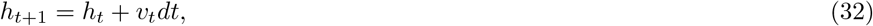

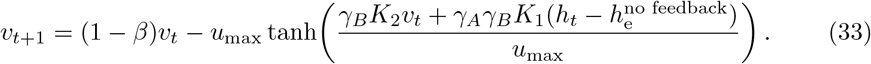

##### Equilibrium and final heading

The equilibrium of the nonlinear no-feedback system satisfies

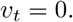

Substituting *v*_*t*_ = 0 into the velocity equation gives

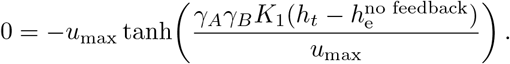

Because tanh(*z*) = 0 only when *z* = 0, the equilibrium heading is

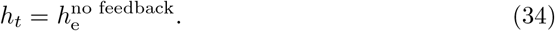

Therefore, the equilibrium of the nonlinear no-feedback model is

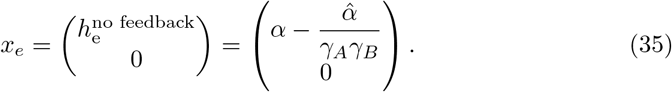

Thus, the saturating nonlinearity does not change the no-feedback equilibrium, but only changes the trajectory used to approach that equilibrium. As in the linear model, the final heading depends on the internal motion gains *γ*_*A*_ and *γ*_*B*_, but not on the control gains *K*_1_ and *K*_2_ or on the acceleration bound *u*_max_.

##### Phase space

The nullclines for the nonlinear model are as follows. For heading we have, as before,

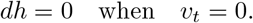

For velocity, we have *dv* = 0 when

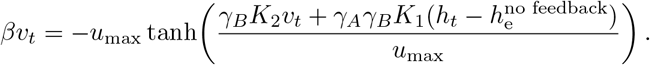

This latter expression cannot be solved analytically but can be solved numerically.

**Note**. There is a distance at which the model switches from being at *v*_term_ to abruptly slowing down. This distance can be approximated by considering where a tangent to the *v*-nullcline intersects *v*_term_. In that region, the dynamics return to the linear regime.

##### Scaling law

The nonlinear model does not obey the scaling law of the linear model. In the linear model, multiplying the starting angle *α* by a constant multiplies the whole trajectory by the same constant. This is because the control signal is linear in the estimated state:

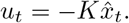

In the nonlinear model,

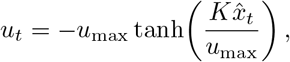

so scaling the initial state does not scale the control signal proportionally. Once the argument of the tanh function becomes large, the control signal approaches ±*u*_max_ regardless of further increases in distance. As a result, trajectories from different starting angles no longer collapse after normalization by *α*.

This failure of the scaling law is a desirable feature of the nonlinear model: it prevents unrealistically large velocities and accelerations while preserving the same equilibrium heading prediction as the linear model.

##### Feedback condition

The no-feedback transformation still holds before and after the feedback update, so the solution for the equilibrium heading remains unchanged. However, the saturating nonlinearity changes how the prediction error affects the immediate control response.

At the feedback time *t*_*fb*_, the visual prediction error is

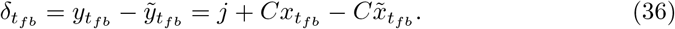

The internal state estimate is then updated according to

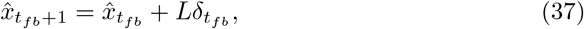

where 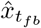 is the estimate immediately before feedback and 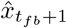 is the estimate immediately after feedback.

The nonlinear control signal immediately before feedback is

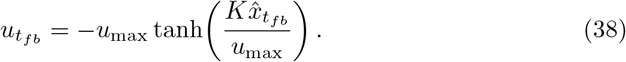

Immediately after feedback, the control signal becomes

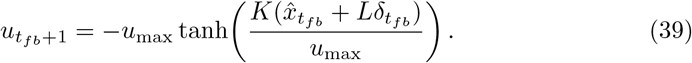

Therefore, the immediate feedback-induced change in control is

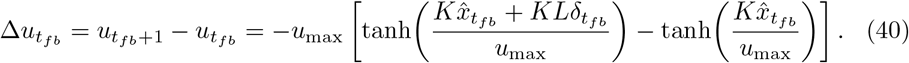

For small prediction errors, this can be approximated by a first-order Taylor expansion:

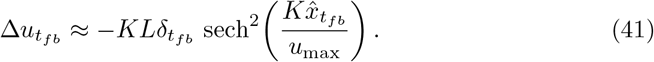

Equation 41 shows that the nonlinear model predicts a state-dependent sensitivity to prediction error. When the current control command is far from saturation,

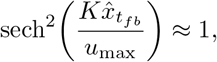

and the feedback response is approximately linear:

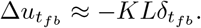

However, when the control input is saturated,

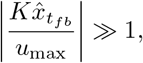

then

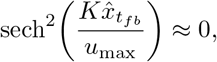

and the same prediction error produces a much smaller change in control.

Thus, unlike the linear model, the nonlinear model predicts that the behavioral effect of prediction error depends on when feedback is delivered during the movement. Prediction errors presented while the controller is saturated should produce relatively small immediate changes in acceleration, whereas prediction errors presented near the deceleration phase, when the controller has returned to the approximately linear regime, should produce larger changes in acceleration.

##### Final heading under feedback

Because the no-feedback transformation still holds after the feedback update, the equilibrium heading under feedback is the same as in the linear model:

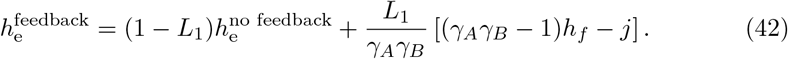

Therefore, the saturating nonlinearity changes the time course of the response to feedback but not the equilibrium heading predicted from the updated internal state. The equilibrium heading depends on the sensory prediction error, the feedback gain *L*_1_, and the internal motion gains *γ*_*A*_ and *γ*_*B*_, but not directly on *K*_1_, *K*_2_, or *u*_max_.

## 5 Data availability

The data used in this study will be made publicly available on the Open Science Framework (OSF).

## 6 Code availability

Custom MATLAB code used for analyses is publicly available on GitHub at https://github.com/yinqihuang/VR_Control.

## Author Contributions

Yinqi Huang: Conceptualization, Methodology, Software, Validation, Formal analysis, Investigation, Project administration, Data curation, Visualization, Writing – original draft, Writing – review & editing, Funding acquisition.

Abhilasha Vishwanath: Methodology, Software, Investigation, Project administration.

Yu Karen Du: Software, Methodology, Data curation.

Matthew F. Watson: Software, Methodology, Investigation.

Osama Asiri: Investigation, Data curation.

Karlee Dakin: Investigation, Data curation.

Donnette Markham: Investigation, Data curation.

Arne D. Ekstrom: Conceptualization, Methodology, Software, Resources, Project administration, Supervision, Funding acquisition, Writing – review & editing.

Robert C. Wilson: Conceptualization, Methodology, Software, Formal analysis, Project administration, Supervision, Funding acquisition, Writing – original draft, Writing – review & editing.

## References

[1] Mittelstaedt, M.-L., Mittelstaedt, H.: Homing by path integration in a mammal. Naturwissenschaften 67(11), 566–567 (1980) 10.1007/BF00450672. Accessed 2026-02-22

[2] Butler, W.N., Smith, K.S., Meer, M.A.A., Taube, J.S.: The head-direction signal plays a functional role as a neural compass during navigation. Current biology: CB 27(9), 1259–1267 (2017) 10.1016/j.cub.2017.03.033

[3] Anastasiou, C., Baumann, O., Yamamoto, N.: Does path integration contribute to human navigation in large-scale space? Psychonomic Bulletin & Review 30(3), 822–842 (2023) 10.3758/s13423-022-02216-8. Accessed 2026-02-22

[4] Taube, J.S.: The head direction signal: origins and sensory-motor integration. Annual Review of Neuroscience 30(1), 181–207 (2007) 10.1146/annurev.neuro.29.051605.112854. Accessed 2026-03-30

[5] Angelaki, D.E., Cullen, K.E.: Vestibular system: the many facets of a multimodal sense. Annual Review of Neuroscience 31, 125–150 (2008) 10.1146/annurev.neuro.31.060407.125555

[6] Arthur, J.C., Philbeck, J.W., Chichka, D.: Non-sensory inputs to angular path integration. Journal of vestibular research : equilibrium & orientation 19(3-4), 111–125 (2009) 10.3233/VES-2009-0354. Accessed 2026-02-22

[7] Angelaki, D.E., Laurens, J.: The head direction cell network: attractor dynamics, integration within the navigation system, and three-dimensional properties. Current Opinion in Neurobiology 60, 136–144 (2020) 10.1016/j.conb.2019.12.002. Accessed 2026-02-22

[8] Kearns, M.J., Warren, W.H., Duchon, A.P., Tarr, M.J.: Path integration from optic flow and body senses in a homing task. Perception 31(3), 349–374 (2002) 10.1068/p3311

[9] Tcheang, L., Bulthoff, H.H., Burgess, N.: Visual influence on path integration in darkness indicates a multimodal representation of large-scale space. Proceedings of the National Academy of Sciences of the United States of America 108(3), 1152–1157 (2011) 10.1073/pnas.1011843108. Accessed 2026-02-22

[10] Etienne, A.S., Jeffery, K.J.: Path integration in mammals. Hippocampus 14(2), 180–192 (2004) 10.1002/hipo.10173

[11] Zhao, M., Warren, W.H.: How you get there from here: interaction of visual landmarks and path integration in human navigation. Psychological Science 26(6), 915–924 (2015) 10.1177/0956797615574952

[12] Zhang, K.: Representation of spatial orientation by the intrinsic dynamics of the head-direction cell ensemble: a theory. The Journal of Neuroscience 16(6), 2112–2126 (1996) 10.1523/JNEUROSCI.16-06-02112.1996. Accessed 2026-03-30

[13] Wilson, R.C., Finkel, L.H.: A neural implementation of the kalman filter. In: Advances in Neural Information Processing Systems 22, pp. 1713–1720. Curran Associates, Inc., Red Hook, NY (2009). Accessed: 2026-03-30. https://proceedings.neurips.cc/paperfiles/…

[14] Petzschner, F.H., Glasauer, S., Stephan, K.E.: A Bayesian perspective on magnitude estimation. Trends in Cognitive Sciences 19(5), 285–293 (2015) 10.1016/j.tics.2015.03.002. Accessed 2026-02-22

[15] Harootonian, S.K., Ekstrom, A.D., Wilson, R.C.: Combination and competition between path integration and landmark navigation in the estimation of heading direction. PLOS Computational Biology 18(2), 1009222 (2022) 10.1371/journal.pcbi.1009222. Accessed 2026-02-22

[16] Kutschireiter, A., Basnak, M.A., Wilson, R.I., Drugowitsch, J.: Bayesian inference in ring attractor networks. Proceedings of the National Academy of Sciences 120(9), 2210622120 (2023) 10.1073/pnas.2210622120. Accessed 2026-03-29

[17] Chen, Y., Zhang, L., Chen, H., Sun, X., Peng, J.: Synaptic ring attractor: A unified framework for attractor dynamics and multiple cues integration. Heliyon 10(16), 35458 (2024) 10.1016/j.heliyon.2024.e35458. Accessed 2026-02-22

[18] Vishwanath, A., Watson, M.F., Gin, M.K., Markham, D.C., Huang, Y., Du, Y.K., Ekstrom, A., Wilson, R.C.: Bayesian cue combination best predicts straightline distance estimation with translated visual landmarks. Neuropsychologia 222, 109351 (2026) 10.1016/j.neuropsychologia.2025.109351. Accessed 2026-03-30

[19] Kessler, F., Frankenstein, J., Rothkopf, C.A.: Human navigation strategies and their errors result from dynamic interactions of spatial uncertainties. Nature Communications 15(1), 5677 (2024) 10.1038/s41467-024-49722-y. Accessed 2026-02-22

[20] Wolpert, D.M., Ghahramani, Z., Jordan, M.I.: An internal model for sensorimotor integration. Science 269(5232), 1880–1882 (1995) 10.1126/science.7569931. Accessed 2026-04-18

[21] Dallos, P., Jones, R.: Learning behavior of the eye fixation control system. IEEE Transactions on Automatic Control 8(3), 218–227 (1963) 10.1109/TAC.1963.1105574. Accessed 2026-04-18

[22] Broucke, M.E.: Adaptive internal model theory of the oculomotor system and the cerebellum. IEEE Transactions on Automatic Control 66(11), 5444–5450 (2021) 10.1109/TAC.2020.3046574. Accessed 2026-04-18

[23] Todorov, E., Jordan, M.I.: Optimal feedback control as a theory of motor coordination. Nature Neuroscience 5(11), 1226–1235 (2002) 10.1038/nn963. Accessed 2026-02-22

[24] Flash, T., Hogan, N.: The coordination of arm movements: an experimentally confirmed mathematical model. The Journal of Neuroscience 5(7), 1688–1703 (1985) 10.1523/JNEUROSCI.05-07-01688.1985. Accessed 2026-03-29

[25] Jagacinski, R.J., Flach, J.M.: Control Theory for Humans: Quantitative Approaches to Modeling Performance, transferred to digital print edn. Erlbaum, Mahwah, NJ (2009)

[26] Franklin, D., Wolpert, D.: Computational mechanisms of sensorimotor control. Neuron 72(3), 425–442 (2011) 10.1016/j.neuron.2011.10.006. Accessed 2026-03-29

[27] Brown, G.L., Seethapathi, N., Srinivasan, M.: A unified energy-optimality criterion predicts human navigation paths and speeds. Proceedings of the National Academy of Sciences 118(29), 2020327118 (2021) 10.1073/pnas.2020327118. Accessed 2026-02-22

[28] Crassidis, J.L., Junkins, J.L.: Optimal Estimation of Dynamic Systems, 2. ed edn. Chapman & Hall CRC applied mathematics and nonlinear science series, vol. 24. CRC Press, Boca Raton, Fla. (2012)

[29] Souman, J.L., Frissen, I., Sreenivasa, M.N., Ernst, M.O.: Walking straight into circles. Current biology: CB 19(18), 1538–1542 (2009) 10.1016/j.cub.2009.07.053

[30] Huang, Y.: Understanding as designed projection. Computational Brain & Behavior (2026) 10.1007/s42113-026-00299-3. Accessed 2026-04-15

[31] Chen, X., McNamara, T.P., Kelly, J.W., Wolbers, T.: Cue combination in human spatial navigation. Cognitive Psychology 95, 105–144 (2017) 10.1016/j.cogpsych.2017.04.003. Accessed 2026-03-30

[32] Stavropoulos, A., Lakshminarasimhan, K.J., Laurens, J., Pitkow, X., Angelaki, D.E.: Influence of sensory modality and control dynamics on human path integration. eLife 11, 63405 (2022) 10.7554/eLife.63405. Accessed 2026-02-22

[33] Cowan, N.J., Ankarali, M.M., Dyhr, J.P., Madhav, M.S., Roth, E., Sefati, S., Sponberg, S., Stamper, S.A., Fortune, E.S., Daniel, T.L.: Feedback control as a framework for understanding tradeoffs in biology. Integrative and Comparative Biology 54(2), 223–237 (2014) 10.1093/icb/icu050. Accessed 2026-05-30

[34] Park, I.M., Vermani, A., Polavieja, G.G.d., Gallego, J., Esfahany, K., Saxena, S., Orger, M., Ijspeert, A., Dowling, M., McNamee, D., Turaga, S.C., Mainen, Z., Paton, J.J., Renart, A.: Integrative neurocybernetic modeling in the era of large-scale neuroscience. arXiv. arXiv:2604.23903 (2026). 10.48550/arXiv.2604.23903. http://arxiv.org/abs/2604.23903 Accessed 2026-05-30

[35] Starrett, M.J., McAvan, A.S., Huffman, D.J., Stokes, J.D., Kyle, C.T., Smuda, D.N., Kolarik, B.S., Laczko, J., Ekstrom, A.D.: Landmarks: A solution for spatial navigation and memory experiments in virtual reality. Behavior Research Methods 53(3), 1046–1059 (2021) 10.3758/s13428-020-01481-6. Accessed 2026-02-22

[36] Huang, Y.: Leveraging virtual reality to understand human spatial navigation. Nature Reviews Psychology 2(11), 662–662 (2023) 10.1038/s44159-023-00243-3. Accessed 2026-03-25

[37] Khalil, H.K.: Nonlinear systems. Hauptbd., 3. ed edn. Prentice Hall, Upper Saddle River, NJ (2002)

